# Proteomic portraits reveal evolutionarily conserved and divergent responses to spinal cord injury

**DOI:** 10.1101/2021.01.27.428528

**Authors:** Michael A. Skinnider, Jason Rogalski, Seth Tigchelaar, Neda Manouchehri, Anna Prudova, Angela M. Jackson, Karina Nielsen, Jaihyun Jeong, Shalini Chaudhary, Katelyn Shortt, Ylonna Gallagher-Kurtzke, Kitty So, Allan Fong, Rishab Gupta, Elena B. Okon, Michael A. Rizzuto, Kevin Dong, Femke Streijger, Lise Belanger, Leanna Ritchie, Angela Tsang, Sean Christie, Jean-Marc Mac-Thiong, Christopher Bailey, Tamir Ailon, Raphaele Charest-Morin, Nicholas Dea, Jefferson R. Wilson, Sanjay Dhall, Scott Paquette, John Street, Charles G. Fisher, Marcel F. Dvorak, Casey Shannon, Christoph Borchers, Robert Balshaw, Leonard J. Foster, Brian K. Kwon

## Abstract

Despite the emergence of promising therapeutic approaches in preclinical studies, the failure of large-scale clinical trials leaves clinicians without effective treatments for acute spinal cord injury (SCI). These trials are hindered by their reliance on detailed neurological examinations to establish outcomes, which inflate the time and resources required for completion. Moreover, therapeutic development takes place in animal models whose relevance to human injury remains unclear. Here, we address these challenges through targeted proteomic analyses of CSF and serum samples from 111 acute SCI patients and, in parallel, a large animal (porcine) model of SCI. We develop protein biomarkers of injury severity and recovery, including a prognostic model of neurological improvement at six months with an AUC of 0.91, and validate these in an independent cohort. Through cross-species proteomic analyses, we dissect evolutionarily conserved and divergent aspects of the SCI response, and establish the CSF abundance of glial fibrillary acidic protein (GFAP) as a biochemical outcome measure in both humans and pigs. Our work opens up new avenues to catalyze translation by facilitating the evaluation of novel SCI therapies, while also providing a resource from which to direct future preclinical efforts.

---

Each year, tens of thousands of individuals suffer an acute traumatic spinal cord injury (SCI) (*1*). The impact of the resulting neurological impairment on the physical, social, and vocational well-being of these individuals is profound (*2*). The burden on health care systems is likewise enormous (*3*). Decades of research in this field have generated numerous therapeutic interventions that have shown promise in preclinical studies (*4, 5*). However, among the handful that have emerged from the laboratory to be tested in the clinic, none have succeeded in demonstrating convincing neurological benefit.

In SCI, the primary traumatic insult to the spinal cord initiates a progressive cascade of secondary injury mechanisms, characterized by ischemia, excitotoxicity, apoptosis, demyelination, inflammatory cell infiltration and cytokine release, and the formation of a glial scar (*6*). Therapeutic approaches generally seek to curtail these responses, in order to improve neurological outcomes via neuroprotection. Interrogation of these processes in human patients is limited by the inaccessibility of spinal cord tissue, and consequently, efforts to unravel this cascade rely almost entirely on animal models (*7*). The underlying assumption is that the pathological responses targeted in the animal setting will be similarly modulated in the injured human spinal cord. However, the failure of efforts to translate promising treatments from animal models into the clinic raises the possibility that important biological differences exist. As such, the dearth of data describing the pathobiology of SCI in human patients represents an obstacle to the development of new therapeutic approaches.

Once a promising intervention for SCI has emerged from the laboratory, the process of establishing efficacy in human patients is exceedingly challenging. Only a handful of large-scale clinical trials in acute SCI have ever been completed, each of which has highlighted the time and tremendous resources necessary for completion (*8*); more recent trials have experienced significant challenges in recruitment, leading to their premature termination. At the heart of this difficulty is the singular reliance on standardized neurological assessments for patient enrollment and evaluation of efficacy. These detailed examinations are challenging or impossible to perform in a substantial fraction of acutely injured patients, due to concomitant trauma, intoxication, or sedation (*9*). Without a baseline evaluation, such individuals are rendered ineligible for clinical trials, severely limiting the pool of recruitable patients. Further compounding this challenge is the considerable variability in spontaneous recovery, even among patients classified similarly by the initial examination, which forces investigators to enroll large numbers of patients in order to discern treatment effects (*10*). With many promising new treatments vying for translation, the inability to validate them clinically presents a bottleneck to the progression of potentially effective therapies.

Clearly, for the field to move forward, new approaches are needed in the evaluation of novel therapeutics. Objective neurochemical biomarkers could catalyze a fundamental shift in the design and conduct of clinical trials for acute SCI (*11*). Diagnostic biomarkers of injury severity could facilitate the enrollment of acutely injured patients unable to complete a detailed neurological examination. Prognostic biomarkers of spontaneous neurological recovery could arguably have even greater utility, by reducing the number of patients needed to adequately power clinical trials. Markers that are similarly modulated in animal models, where outcome measures currently used to assess therapeutic efficacy lack a direct correlate in human, could provide a basis to prioritize therapies for translation into human trials (*12*).

Here, we describe a targeted proteomic analysis of CSF and serum samples serially collected over the first five days following acute SCI in a population of 111 patients. In parallel, we characterized CSF and serum samples from a porcine model of SCI, using an identical proteomic approach. In the patient population, we mine this resource to develop single-and multi-protein biomarkers of baseline injury severity and neurological recovery at six months post-injury, and validate these in an independent cohort, demonstrating highly accurate patient stratification and prognostication. Through comparative proteomic analyses, we establish both conserved biological outcomes and species-specific responses between human SCI and a large animal model, including the discovery of glial fibrillary acidic protein (GFAP) as a cross-species outcome measure. With a total of 1,329 samples analyzed, this biobank defines an unprecedented opportunity to interrogate the biology of acute traumatic SCI.

## Results

### Overview of study cohorts

A total of 111 individuals sustaining an acute SCI were recruited through a multicenter prospective observational trial. Patients underwent a neurological examination at baseline, and again at six months post-injury to determine the extent of neurological improvement. Clinical characteristics are provided in **Table S1**. CSF and serum samples were collected serially over the first five days post-injury from all patients, and were additionally obtained from 21 uninjured patients who served as negative controls. In total, 910 samples were collected and subjected to proteomic profiling, including 450 from the CSF and 460 from the serum, with between 63-100 samples from each biofluid collected at each timepoint (**Fig. 1A**).

**Fig. 1:**
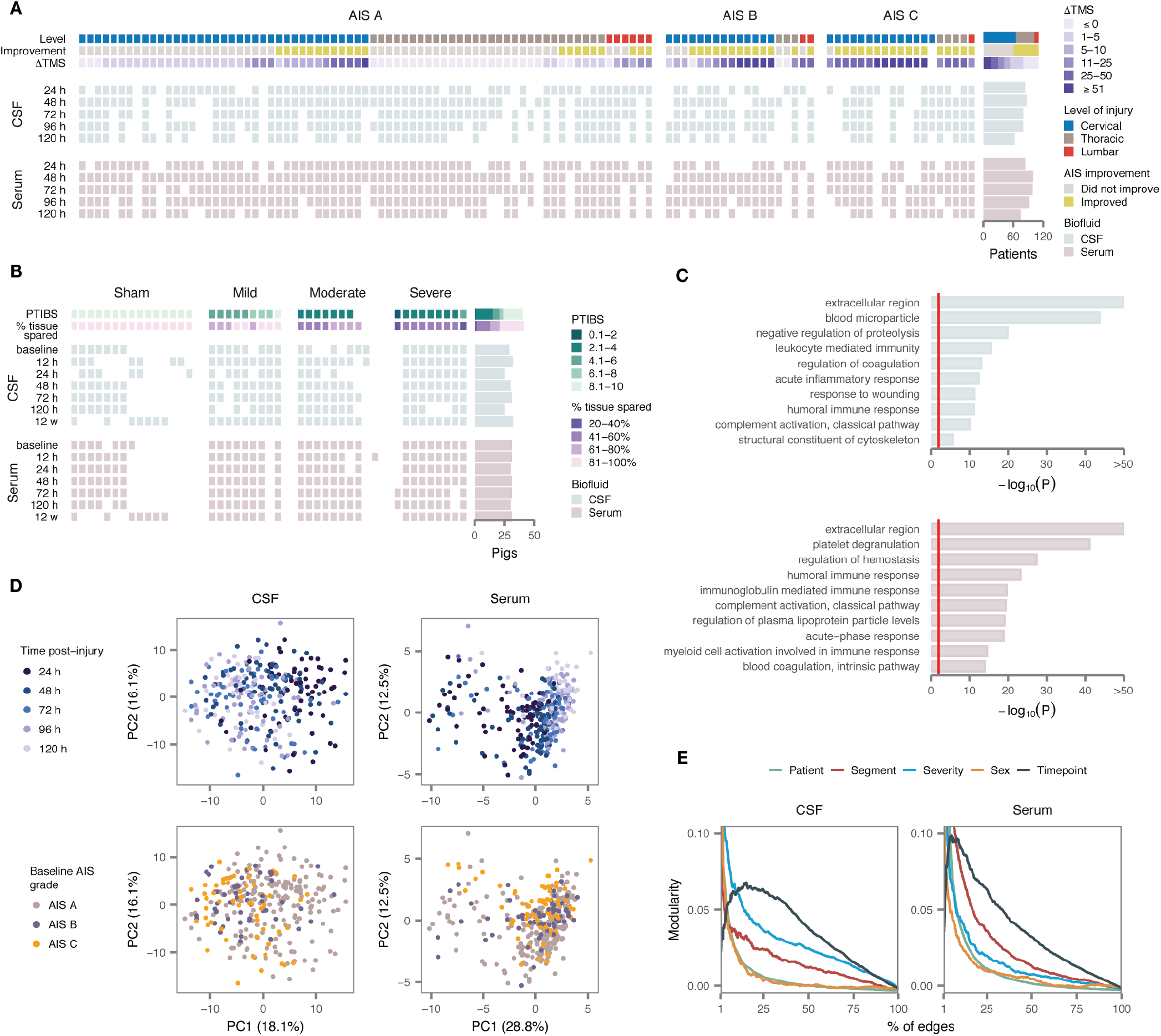
Targeted proteomic profiles of CSF and serum in human and pig SCI. (A) Overview of the human cohort (*n* = 111 patients), including key clinical measures (level of injury, AIS conversion, and change in TMS at six months post-injury), top, and sample collection at each of five timepoints in the CSF, middle, and serum, bottom. (B) Overview of the pig cohort (*n* = 43 animals), including key experimental outcomes (PTIBS score at 12 weeks post-injury, percentage of histological tissue sparing), top, and sample collection at each of seven timepoints in the CSF, middle, and serum, bottom. (C) GO enrichment analysis of proteins targeted by synthetic peptides in the CSF, top, and serum, bottom. Red line shows 1% FDR. (D) PCA of human CSF and serum proteomes, with samples colored by time post-injury, top, or baseline AIS grade, bottom. (E) Modularity analysis of CSF and serum proteomes with samples grouped by patient, level of injury (cervical, thoracic, or lumbar spinal cord), injury severity (baseline AIS grade), sex, or time post-injury. Modularity is shown as a function of the number of edges between samples used to construct the network, as a percentage of the total number of possible edges.

In parallel, we conducted an experimental study in our previously described porcine model of traumatic SCI (*13*). The pig was selected on the basis of its anatomical and immunological similarities to human (*14*), in particular with respect to the dimensions of the spinal canal, which permit serial CSF collection through indwelling intrathecal ports (**Fig. 1A-B**) (*15*). A total of 28 pigs received contusion SCIs of three different severities (mild, moderate, or severe; 9-10 pigs per group), and an additional 15 pigs received sham laminectomies (**Fig. 1B**). CSF and serum samples were collected 15 minutes prior to injury, and serially over six timepoints between 12 h and 12 weeks post-injury. Hindlimb neurological recovery was assessed weekly in all animals until 12 weeks, at which point animals were euthanized and spinal cords were collected for histological analysis (**Fig. S1C-H**). In total, 419 samples were profiled from the pig study, including 204 from the CSF and 215 from the serum.

### Development of targeted proteomic assays for CSF and serum

Accurate measurement of protein abundance in the CSF and serum is highly challenging, owing to the complexity and dynamic range of protein concentrations in these biofluids. In order to achieve sensitive and accurate protein quantitation, we designed targeted mass spectrometric assays based on multiple reaction monitoring (MRM) in the serum and parallel reaction monitoring (PRM) in the CSF. Target proteins were selected on the basis of an extensive literature review, in conjunction with discovery proteomic analyses of human CSF. We subsequently developed new PRM assays for 334 peptides, targeting 325 distinct CSF proteins, which in combination with 270 previously described MRM assays (*16, 17*) afforded coverage of 491 proteins (**Table S2**). Gene Ontology (GO) terms enriched among proteins targeted by synthetic peptides reflected immunological and inflammatory pathways, cytoskeletal and secreted proteins, and coagulation (**Fig. 1C**). Isotopically labeled synthetic peptides were spiked into all samples at defined concentrations, allowing direct comparison of protein abundance across the study co-hort. The complete dataset of CSF and serum protein abundance data from both species is provided in **Table S3**.

### Proteomic portraits of SCI across tissues and species

To obtain a global overview of the dataset, we first performed principal component analysis (PCA) of the human cohort. Samples separated on the first principal component by time-point, rather than clinical severity, in both biofluids, suggesting that time postinjury is the primary driver of variation in protein abundance following acute SCI (**Fig. 1D** and **Fig. S2A-B**). Similar trends were apparent in analyses of the pig samples (**Fig. S2C-D**).

To more formally quantify these trends, we constructed networks in both the serum and CSF, in which nodes corresponded to samples and edges were drawn between pairs of samples with correlated protein abundance, and carried out a modularity analysis of each network (*18*). Grouping samples by timepoint consistently yielded higher modularity than grouping by injury severity, confirming the trends observed visually by PCA (**Fig. 1E**). Interestingly, in the serum, grouping by the injured spinal segment also yielded high modularity, suggesting the neurological level of injury shapes patterns of serum protein abundance within timepoints.

### Profound proteomic alterations in acute SCI

To date, interrogation of the molecular alterations that follow acute SCI has relied almost entirely on rodent models (*19*). Our large patient cohort provides an opportunity to define changes in the CSF and serum proteome of human SCI patients at a scale that has not previously been possible. We therefore first sought to characterize differential protein abundance following acute SCI (**Table S4**).

In the CSF, SCI caused profound changes in protein abundance. At 24 h post-injury, 39% of targeted proteins displayed statistically significant differences relative to uninjured controls, at a 5% false discovery rate (FDR; **Fig. 2A**, left, and **Fig. S3A**). Similar proportions of proteins were differentially expressed (DE) between 48 h and 120 h (**Fig. 2A**, right). Many displayed dramatic alterations; for instance, the abundance of GFAP in the CSF differed by 25-fold between SCI patients and controls (**Fig. 2B**).

**Fig. 2:**
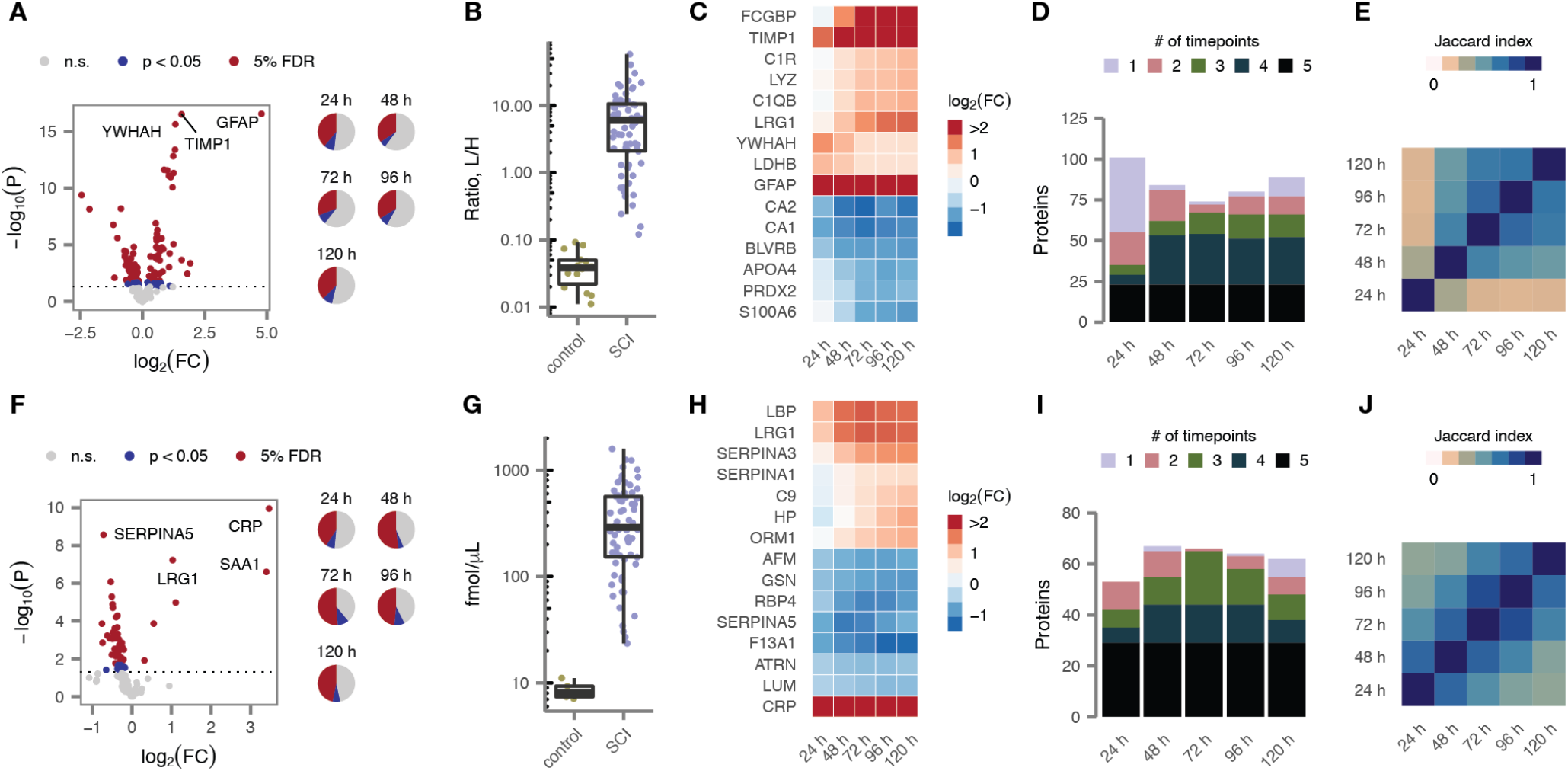
Profound alterations of the CSF and serum proteomes in acute SCI. (A) Volcano plot of differential protein abundance at 24 h post-injury in CSF samples from patients with acute SCI compared to uninjured controls. (B) Abundance of GFAP in CSF samples from acute SCI patients and uninjured controls. (C) Time courses of differential protein abundance (log-fold change, relative to uninjured controls) over the first five days post-injury for fifteen of the most profoundly altered CSF proteins. (D) Number of differentially expressed CSF proteins at each of the first five days post-injury in acute SCI patients, grouped by the number of timepoints at which each protein was found to be differentially expressed. (E) Similarity of proteome alterations between all possible pairs of timepoints, as quantified by the Jaccard index between the differentially expressed proteins at each timepoint. (F) As in (A), but showing differential protein abundance in the serum at 24 h post-injury. (G) Abundance of CRP in serum samples from acute SCI patients and uninjured controls. (H-J) As in (C-E), but showing serum proteins.

Although a handful of the most significant associations were reproduced across all five timepoints, DE proteins displayed varying temporal patterns of up-or downregulation (**Fig. 2C** and **Fig. S3B**). For instance, GFAP was maximally upregulated immediately after injury, whereas complement proteins exhibited a delayed increase in abundance. Moreover, although most DE proteins were identified at a minimum of two adjacent timepoints, a substantial proportion were uniquely identified within 24 h of the initial injury (**Fig. 2D**). Using the Jaccard index to quantify the overlap in DE proteins over each of the first five days post-injury, the 24 h timepoint emerged as a clear outlier (**Fig. 2E**). These observations suggest that a distinct set of pathophysiological processes are active in the spinal cord parenchyma during the most acute phase of the response to SCI.

Similarly marked changes were observed in the serum proteome after SCI, with 41-52% of proteins DE in SCI patients over the five-day time course (**Fig. 2F** and **Fig. S3C**). Acute-phase reactants such as C-reactive protein (CRP; **Fig. 2G**) were among the most strongly altered proteins. However, a less dramatic temporal course was apparent in the serum (**Fig. 2H** and **Fig. S3D**): most DE proteins were shared between adjacent timepoints, and the 24 h timepoint was not an obvious outlier (**Fig. 2I-J**). These results suggest the serum proteome reflects systemic responses to SCI that develop over longer timescales than those occurring within the central nervous system itself.

A unique aspect of our dataset is its time course design, whereby samples were collected serially from acute SCI patients over the first five days post-injury. In univariate analyses, nearly all proteins exhibited significant differences in abundance over this time course (**Fig. S3E**). To better understand the pathophysiological responses that are activated over the first five days after acute human SCI, we applied *k*-means consensus clustering (*20*) to identify modules of temporally co-regulated proteins. Seven such modules were identified in each biofluid (**Fig. S4A-D**). These included groups of proteins that were up- or down-regulated immediately after injury but had returned to baseline by five days, as well as groups that displayed more delayed patterns of up- and down-regulation, or which were up- or downregulated for the entire time course (**Fig. S4E-H**). Analysis of enriched GO terms suggested functional correlates of each module (**Fig. S4I-J**). Taken together, these analyses provide an initial description of the temporal profile of molecular changes in human CSF and serum following acute SCI.

### Univariate analysis identifies protein biomarkers of SCI severity and recovery

Next, we sought to identify individual proteins whose abundance in the serum or CSF was associated with baseline injury severity, as quantified by the baseline AIS grade, or neurological recovery, as quantified either by the change in total motor score (ΔTMS) or AIS grade improvement at six months post-injury (**Table S5**). We initially focused on samples collected at 24 h. At this timepoint, a total of 23 proteins were associated with baseline AIS grade in the CSF, all of which were more abundant in patients with more severe injuries (**Fig. 3A**); the most statistically significant of these associations was to GFAP (**Fig. 3B**). Fourteen CSF proteins were significantly associated with ΔTMS (**Fig. 3C**), of which eleven were also associated with baseline AIS grade, including GFAP (**Fig. 3D**). After correction for multiple hypothesis testing, no proteins were associated with AIS grade improvement (**Fig. 3E**). In parallel analyses of serum samples collected at 24 h, only a single association was identified, between coagulation factor F11 (F11) and ΔTMS.

**Fig. 3:**
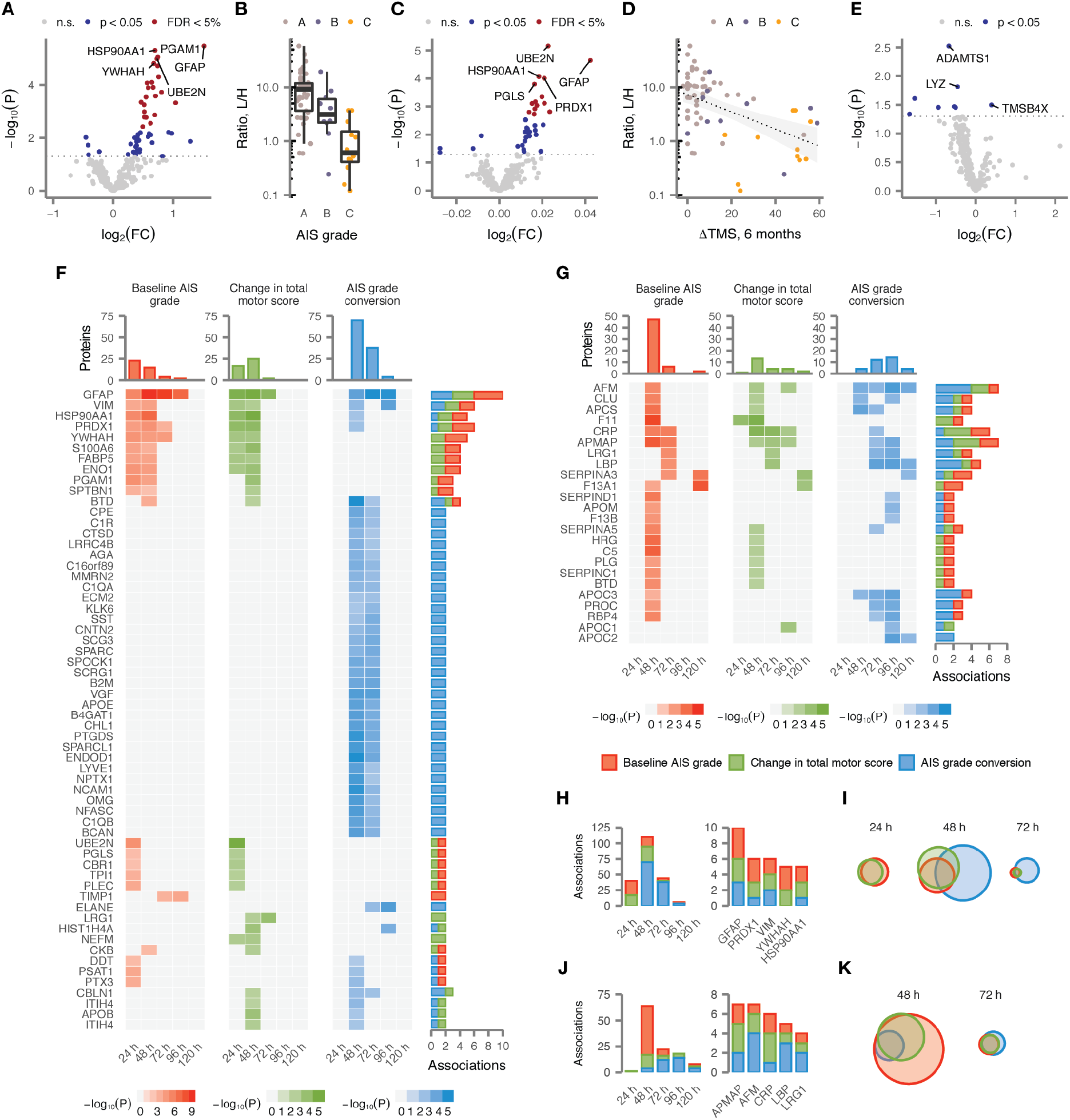
Univariate protein biomarkers of SCI severity and recovery. (A) Volcano plot of differential protein abundance as a function of injury severity, as measured by the baseline AIS grade, in CSF samples at 24 h post-injury. (B) Abundance of GFAP in CSF samples from acute SCI patients, stratified by baseline AIS grade. (C) Volcano plot of differential protein abundance as a function of neurological recovery, as measured by the change in TMS at six months post-injury, in CSF samples at 24 h post-injury. (D) Abundance of GFAP in CSF samples from acute SCI patients, stratified by change in TMS at six months post-injury. (E) Volcano plot of differential protein abundance as a function of neurological recovery, as measured by AIS grade improvement over baseline at six months post-injury, in CSF samples at 24 h post-injury. (F) Statistical significance of associations between CSF protein abundance and three clinical outcomes over the first five days post-injury for 59 proteins with two or more significant associations. Grey squares indicate associations that were not significant after correction for multiple hypothesis testing. (G) As in (F), but for 24 serum proteins with two or more significant associations. (H) Number of statistically significant associations between CSF protein abundance and three clinical outcomes per timepoint, left, and for the five CSF proteins with the most recurrent associations, right. (I) Overlap between CSF proteins significantly associated with three clinical outcomes at 24 h, 48 h, and 72 h post-injury. (J) As in (H), but for serum proteins. (K) As in (I), but for serum proteins at 48 h and 72 h.

Given the dynamic temporal patterns of protein abundance observed in both biofluids, we subsequently repeated these analyses over each timepoint between 48 h and 120 h. In the CSF, a total of 106 proteins were significantly associated with injury severity or neurological recovery within at least one timepoint (**Fig. S5A-C**). We searched specifically for proteins with recurrent associations, spanning multiple timepoints or clinical outcomes, reasoning that these would be substantially less likely to represent spurious hits, and identified a total of 59 proteins with at least two statistically significant associations to injury severity or recovery (**Fig. 3F**). Among these, GFAP emerged as the most robust marker, with ten significant associations across all timepoints (**Fig. 3H**). The number of significant associations peaked at 48 h and subsequently declined, with none identified by 120 h (**Fig. 3H**). Associations to baseline AIS grade and change in TMS largely overlapped, but AIS grade improvement was reflected by a distinct proteomic signature (**Fig. 3I**).

We then carried out a parallel analysis of recurrently associated proteins in the serum (**Fig. S5D-F**). Despite the dearth of associations at 24 h post-injury, the serum proteome was substantially more informative with respect to clinical outcomes at later timepoints. A total of 24 proteins exhibited two or more statistically significant associations (**Fig. 3G**). As in the CSF, the number of associations to clinical outcomes peaked at 48 h, but a subset of serum proteins carried diagnostic or prognostic value as late as 120 h (**Fig. 3J**). Moreover, as in the CSF, there was substantial overlap between proteins associated with neurological recovery or base-line severity (**Fig. 3K**).

Together, these results define dozens of CSF and serum protein biomarkers with recurrent associations to injury severity or recovery over the first five days post-injury, and suggest that protein abundance at 48 h generally carries the greatest prognostic value.

### Machine learning defines highly accurate multi-protein diagnostics

Next, we sought to optimize the accuracy of patient stratification or prognostication by combining diagnostic or prognostic information across multiple proteins simultaneously, through multivariate analyses. To this end, we developed a comprehensive machine-learning pipeline, implementing a total of 22 different classification and regression models. We then applied this pipeline to CSF and serum proteomics data from each timepoint.

We first directed our machine-learning pipeline to develop multi-protein diagnostic models of baseline AIS grade. Models trained on CSF protein concentrations at 24 h or 48 h both achieved good accuracy in fivefold cross-validation (81% and 82%, respectively; **Fig. 4A**). However, the balanced accuracy, which adjusts for differences in the prevalence of each AIS grade, was substantially higher at 48 h (72% vs. 65%; **Fig. S6A**). Closer inspection suggested that both models had difficulty correctly identifying AIS B patients (**Fig. 4B** and **Fig. S6B**). Serum models were generally less accurate (**Fig. 4A**, **Fig. S6A** and **C**). Although the severity of the injury is likely to be clinically apparent by 48 h in individuals able to perform a baseline neurological exam, a substantial proportion of patients remain poorly examinable up to 72 h post-injury, due to intubation and sedation following urgent surgical intervention. These patients would conceivably be candidates for emerging therapies delivered at sub-acute time-points, suggesting a diagnostic tool based on CSF protein abundance at 48 h could have substantial clinical utility.

**Fig. 4:**
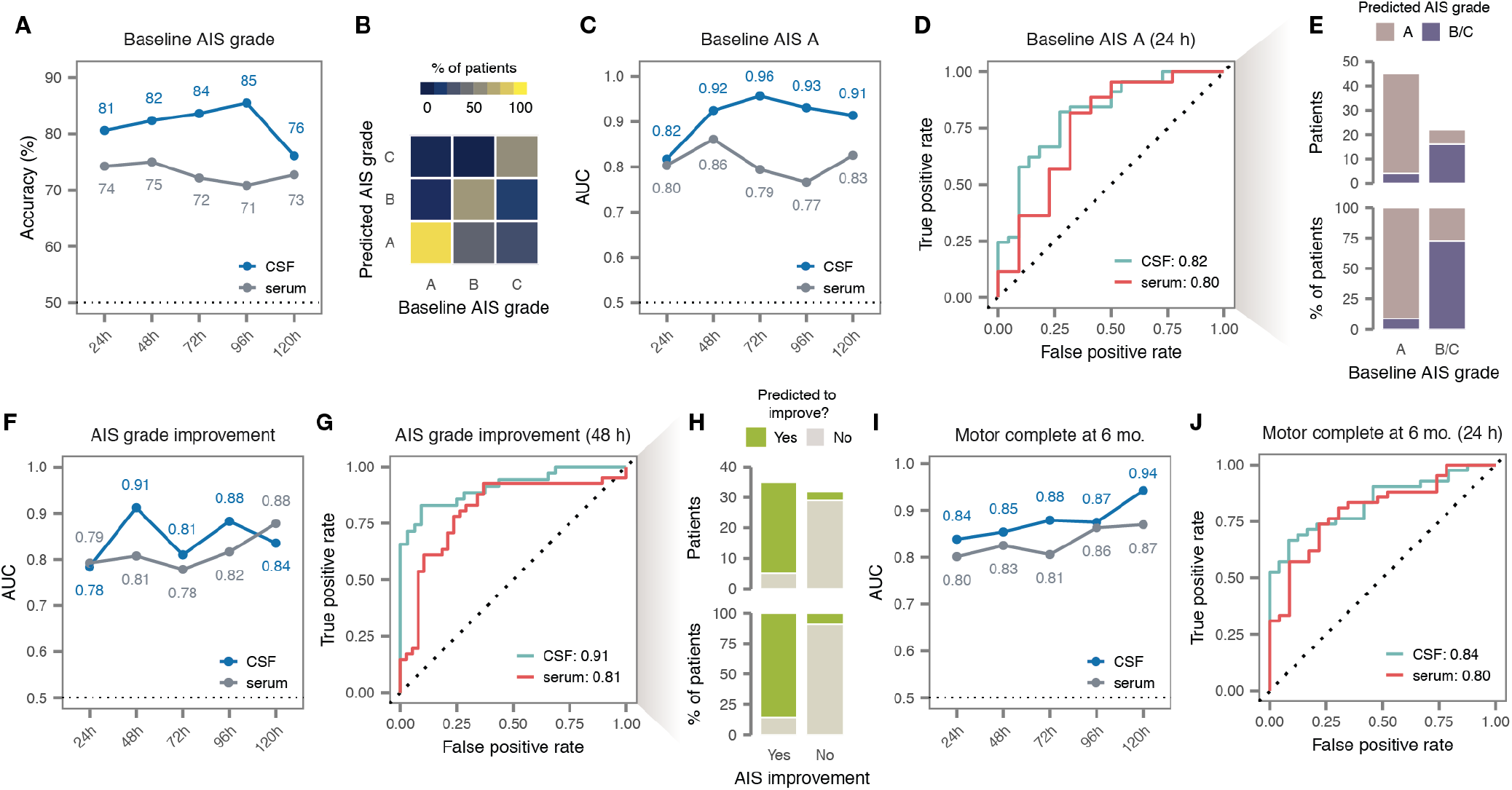
Multivariate protein biomarkers of SCI severity and recovery. (A) Cross-validation accuracy of multivariate diagnostic models trained to stratify patients by baseline AIS grade, at timepoints between 24 and 120 h post-injury. (B) Confusion matrix of the best CSF diagnostic model of baseline AIS grade at 48 h post-injury. (C) Cross-validation AUC of multivariate diagnostic models trained to discern patients with a baseline AIS grade of A, at timepoints between 24 and 120 h post-injury. (D) ROC curves of the best CSF and serum diagnostic models at 24 h post-injury. (E) Predictions made by the best CSF diagnostic model at 24 h post-injury. (F) Cross-validation AUC of multivariate prognostic models trained to predict improvement in AIS grade at six months post-injury, relative to baseline, at timepoints between 24 and 120 h post-injury. (G) ROC curves of the best CSF and serum prognostic models of AIS grade improvement at 48 h post-injury. (H) Predictions made by the best CSF prognostic model at 48 h post-injury. (I) Cross-validation AUC of multivariate prognostic models trained to predict motor complete vs. incomplete injury at six months, at timepoints between 24 and 120 h post-injury. (J) ROC curves of the best CSF and serum prognostic models of motor complete vs. incomplete injury at 24 h post-injury.

Motivated by the observation that our diagnostic models struggled to differentiate intermediate injury severities, we also asked whether we could specifically discriminate AIS A from AIS B or C patients. Indeed, CSF models demonstrated strong performance in identifying AIS A patients as early as 24 h, achieving a maximum AUC of 0.82 and an accuracy of 85% (**Fig. 4C-E** and **Fig. S6D**). Performance was only slightly poorer in the serum (**Fig. 4C-D**). These findings suggest the possibility of an acute diagnostic test to specifically identify the subset of patients in whom the likelihood of spontaneous recovery is lowest.

### Multi-protein biomarkers predict neurological improvement at six months post-injury

Next, we applied the same machine-learning pipeline to develop multi-protein models capable of prognosticating ultimate neurological recovery. With respect to ΔTMS, the best performance was achieved by a model trained on the CSF proteome at 48 h, which achieved a coefficient of determination (r^2^ score) of 0.51 (**Fig. S6E-F**). However, performance was poorer for CSF models at 24 h, and in the serum at either timepoint. Previous studies have defined ΔTMS beyond a prespecified threshold, typically 6-7 points, as a primary outcome (*21, 22*), and consequently we investigated whether better accuracy could be achieved in predicting a ΔTMS of 7 or greater. Indeed, we found that serum models achieved an AUC of 0.83 and an accuracy of 78% at 24 h on this task (**Fig. S6G-I**). Thus, while predicting the exact magnitude of the change in TMS is challenging, predicting whether there will be a clinically meaningful improvement can be achieved with reasonably good accuracy.

We also sought to predict neurological recovery as defined by improvement in the AIS grade, another measure that has widely been used as a primary endpoint in clinical trials (*23, 24*) (**Fig. 4F** and **Fig. S6J**). The best-performing models made use of CSF proteomics data at 48 h post-injury, achieving an exceptionally high AUC of 0.91 and an accuracy of 88% (**Fig. 4G-H**). This performance suggests the possibility of highly accurate identification of patients likely to demonstrate spontaneous neurological improvement, with implications for both the design of clinical trials and acute patient management.

In view of these successes, we further evaluated the possibility of directly prognosticating neurological impairment at six months. Prediction of six-month AIS grade was not substantially better than random expectation (**Fig. S6K**). However, we found that multi-protein models could reliably distinguish motor complete vs. incomplete injuries at six months (that is, a six-month AIS grade of A/B vs. C/D; **Fig. 4I** and **Fig. S6L**). At 24 h, the best serum model achieved an AUC of 0.80 and an accuracy of 80% (**Fig. 4J** and **Fig. S6M**).

Taken together, these results suggest it is possible to predict much of the observed variation in neurological recovery at six months post-injury on the basis of protein abundance in acutely collected CSF and serum samples.

### Validation of single- and multi-protein biomarkers in an independent clinical cohort

Having established both single-and multi-protein biomarkers of SCI severity and recovery, we next sought to validate these in an independent clinical cohort. Samples from the validation cohort (*n* = 20 patients) were processed separately from the discovery cohort and blinded to investigators until after finalization of all statistical analyses. A large fraction of the alterations following acute SCI were reproduced within each biofluid and time-point, and effect sizes were highly correlated between cohorts (r ≥ 0.90; **Fig. 5A-D** and **Fig. S7A-B**). A total of 28 associations to injury severity or recovery were reproduced at 5% FDR, with a further 30 nominally significant (**Fig. 5E-F**). In the CSF, GFAP again stood out as the most promising candidate, with six associations validated at nominal significance or better (**Fig. S7C-E**). In the serum, associations between LRG1 at 72 h and both ΔTMS and AIS grade improvement were reproduced at 5% FDR (**Fig. S7F-G**).

**Fig. 5:**
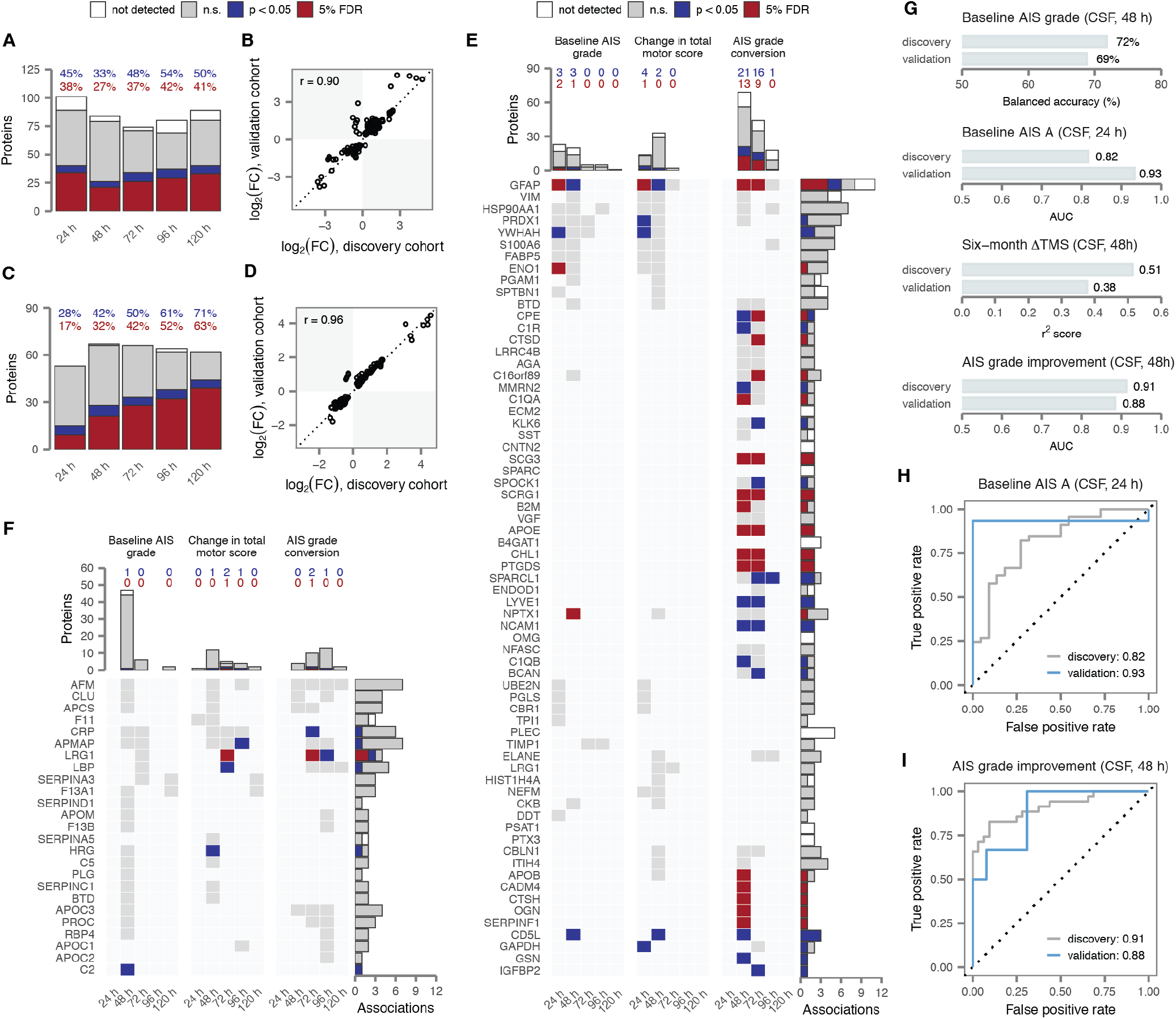
Validation of univariate and multivariate biomarkers in an independent cohort. (A) Proportion of univariate alterations in CSF protein abundance after acute SCI that were replicated at nominal significance or after multiple hypothesis testing correction, were not significant, or were not quantified in the validation cohort. (B) Correlation between log-fold changes between the discovery and validation cohorts for proteins with replicated alterations. (C-D) As in (A-B), but for alterations in serum protein abundance. Cross-validation AUC of multivariate diagnostic models trained to discern patients with a baseline AIS grade of A, at timepoints between 24 and 120 h post-injury. (E) Replicated associations between CSF protein abundance and three clinical outcomes over the first five days post-injury for 59 proteins with two or more significant associations (as shown in **Fig. 3F**) and an additional eight proteins with a single association replicated with at least nominal significance. (F) As in (E), but for the 24 serum proteins with two or more significant associations (as shown in **Fig. 3G**) and one additional protein with a single association replicated with nominal significance. (G) Performance of four preregistered multivariate CSF models in an independent validation cohort. (H) ROC curves of the best CSF diagnostic model of AIS A patients at 24 h post-injury (as shown in **Fig. 4D**) in the discovery and validation cohorts. (I) ROC curves of the best CSF prognostic models of AIS grade improvement at 48 h post-injury (as shown in **Fig. 4G**) in the discovery and validation cohorts.

Among the multivariate models described above, six were selected and preregistered for validation, including four CSF and two serum models. All four CSF models achieved performance comparable to that observed in the discovery cohort (**Fig. 5G**); serum models, in contrast, did not demonstrate convincing accuracy (**Fig. S7H**). Remarkably, in the validation cohort, patients with a baseline AIS grade of A could be identified with an AUC of 0.93 from 24 h CSF (**Fig. 5H**) and AIS grade improvement at six months could be predicted with an AUC of 0.88 from 48 h CSF (**Fig. 5I**). Models of baseline AIS grade and ΔTMS based on 48 h CSF also achieved good performance (**Fig. S7I-J**). Although not among the six preregistered models, we additionally observed that motor complete injuries at six months could be identified from 24 h CSF with an AUC of 0.96 in the validation cohort (**Fig. S7K**).

Thus, through blinded evaluation in an independent co-hort, enrolled and processed after the discovery cohort to emulate a prospective clinical study, we successfully validated dozens of individual proteins and four multi-protein models for patient stratification and prognostication in SCI.

### Proteomic alterations in a large animal model of SCI

The vast majority of our knowledge concerning the pathophysiological responses to acute SCI has emerged from studies in animal models. Through these studies, many therapies have emerged that effectively target the secondary injury responses triggered after SCI in rodents and other species. However, the subsequent failure of these apparently promising agents in human clinical trials raises the possibility that important biological differences exist between animal models and human SCI. The difficulty of translating novel therapies is compounded by the fact that functional or anatomical outcome measures assessed in animal models, such as “hindlimb locomotor function” or “white matter sparing at the injury site”, cannot be measured in humans, or may have limited relation to clinically relevant outcomes. Establishing biochemical markers that are predictive of neurological outcomes in both animal models and human patients could provide endpoints for preclinical studies that are directly informative about human SCI. We therefore sought to leverage our parallel study of SCI in a large animal model in order to identify biochemical markers of injury severity and recovery that could serve as translational linkages between species.

We first a ddressed whether, in acombined a nalysis of human and pig samples, proteomic profiles would segregate globally by species or by time post-injury, the dominant source of within-species variation. To this end, we performed PCA on a combined matrix of human and pig samples at four matching timepoints (24 h, 48 h, 72 h, and 120 h). Reassuringly, samples separated primarily by time post-injury, rather than by species, suggesting that divergences between species are not so pronounced as to preclude extrapolation from one organism to another (**Fig. 6A**). Modularity analysis quantitatively confirmed this trend (**Fig. S8A**). Parallel analyses yielded similar results in the serum (**Fig. S8B-C**).

**Fig. 6:**
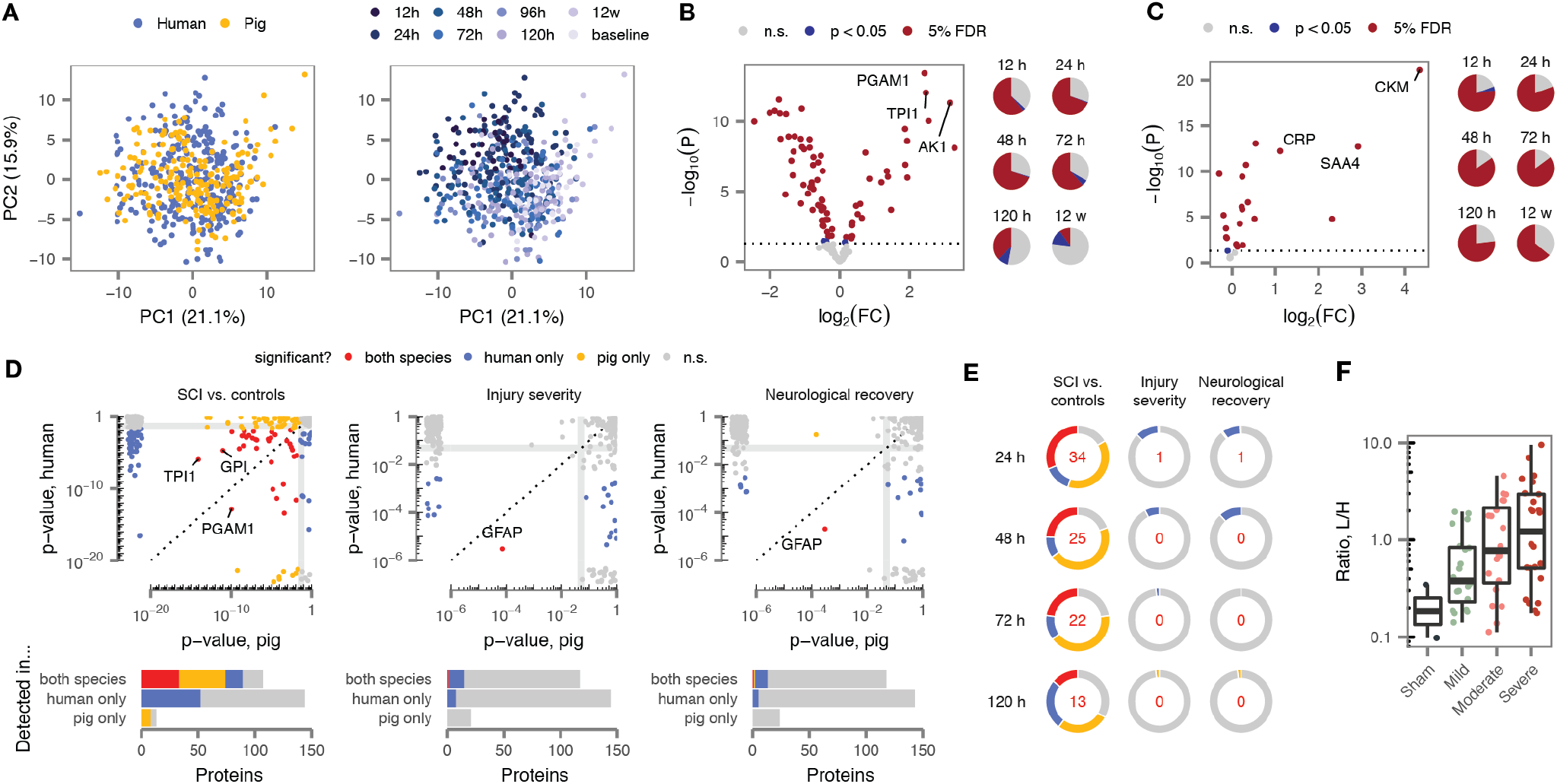
Proteomic profiles of a large animal model define cross-species biomarkers of SCI. (A) Principal component analysis of human and pig CSF proteomes, with samples colored by species, left, or time post-injury, right. (B-C) Volcano plots of differential protein abundance at 24 h post-injury in CSF (B) and serum (C) samples from injured pigs, compared to samples drawn from the same pigs at baseline (15 min prior to injury). Pie charts show the proportion of differentially expressed proteins between 12 h and 12 w post-injury. (D) Top, p-values for differential protein abundance at 24 h in human and pig, in comparisons of individuals with acute SCI and uninjured controls, left; as a function of injury severity, middle; or as a function of neurological recovery, right. Marginal plots show p-values for proteins quantified in human (pig) only. Bottom, number of proteins with statistically significant differential abundance in both species, human only, pig only, or neither, among proteins quantified in both species, human only, or pig only. (E) Proportion of human and pig proteins that are differentially abundant in both species, one species, or neither, at all four matched timepoints following acute SCI, left; as a function of injury severity, middle; or as a function of neurological recovery, right. Inset numbers show proteins that were differentially abundant in both species. (F) Abundance of GFAP in pig CSF samples at 24 h post-injury as a function of injury severity.

Having established a basis for cross-species comparison, we next sought to identify differences in protein abundance between CSF and serum samples collected from injured pigs, and matching samples drawn 15 min prior to injury (**Table S7**). Despite the comparatively lower number of observations in pig (*n* = 25-32 samples per biofluid and timepoint), this repeated-measures design provided excellent statistical power to detect proteomic alterations in SCI, with 37-69% of proteins DE over the first five days post-injury (**Fig. 6B-C** and **Fig. S8D-G**). Several of these changes persisted until 12 weeks post-injury, providing an initial description of proteomic alterations in the chronic phase of SCI. Similarly, we performed univariate analyses of baseline severity, neurological recovery, and histological tissue sparing in the pig CSF and serum, identifying thirteen proteins with recurrent associations to experimental variables in the CSF, and four in the serum (**Fig. S8H-I**). The majority of these associations were detected at 12 h, although a subset were detected over multiple timepoints (**Fig. S8J-K**).

We next took advantage of our parallel univariate analyses in both human and pig to directly compare protein alterations across species, focusing our analysis primarily on the CSF due to the relatively small number of associations in the serum of either species. A substantial fraction of CSF proteins were comparably modulated following SCI in both human and pig, with 14-31% of proteins DE in both species at matching timepoints (**Fig. 6D-E** and **Fig. S9A**). Accounting for incomplete power by using the *π*_1_ statistic to estimate the proportion of null hypotheses (*25*) further increased the estimated degree of overlap, to 57-81% (**Fig. S8L-M**).

These data indicated that the proteomic alterations that accompany acute SCI are broadly conserved between human and pig. In contrast, markers of injury severity or recovery were more species-specific. Across all four matching timepoints, only two associations were reproduced in both species (**Fig. 6E** and **Fig. S9B-D**). Remarkably, these implicated the same protein at the same timepoint: CSF levels of GFAP at 24 h were correlated to both baseline injury severity and hindlimb neurological recovery in pigs (**Fig. 6F** and **Fig. S8N**). This finding immediately suggests a biochemical outcome measure with direct relevance to human injury that could be used to monitor injury progression and therapeutic response in animal studies, with direct relevance to human injury.

### Convergent proteomic signatures in human and porcine SCI

Although univariate analyses successfully identified individual proteins altered in both human and pig SCI, we reasoned that relying on a simple overlap between DE proteins in either species could reduce our sensitivity to detect small but concordant changes. To overcome this limitation, we performed a threshold-free analysis using rank-rank hypergeometric overlap (RRHO) (*26*) to identify convergent proteomic signatures of acute SCI, injury severity, and recovery. We applied this method to compare all pairs of time-points evaluated in either species, reasoning that equivalent responses may not be activated at perfectly matching time-points between species.

We first applied RRHO analysis to compare proteomic signatures of acute SCI (**Fig. 7A**). As anticipated from our univariate analyses, highly significant overlap was observed between approximately matching timepoints (**Fig. 7B**). Surprisingly, however, there was more limited overlap between mismatched timepoints, suggesting substantial dynamic evolution in the molecular response to SCI over the first five days post-injury. The greatest overlap was observed at 24 h and earlier, indicating the most acute phase of the response to SCI is the most strongly conserved between species. Subacute and chronic phases also showed a broad alignment, with significant overlap between the human protein signature of SCI at 120 h and the pig signature between 72 h and 12 w. This observation suggests that many of the molecular alterations that persist into the chronic stage of the injury have been established in human by 120 h post-injury.

**Fig. 7:**
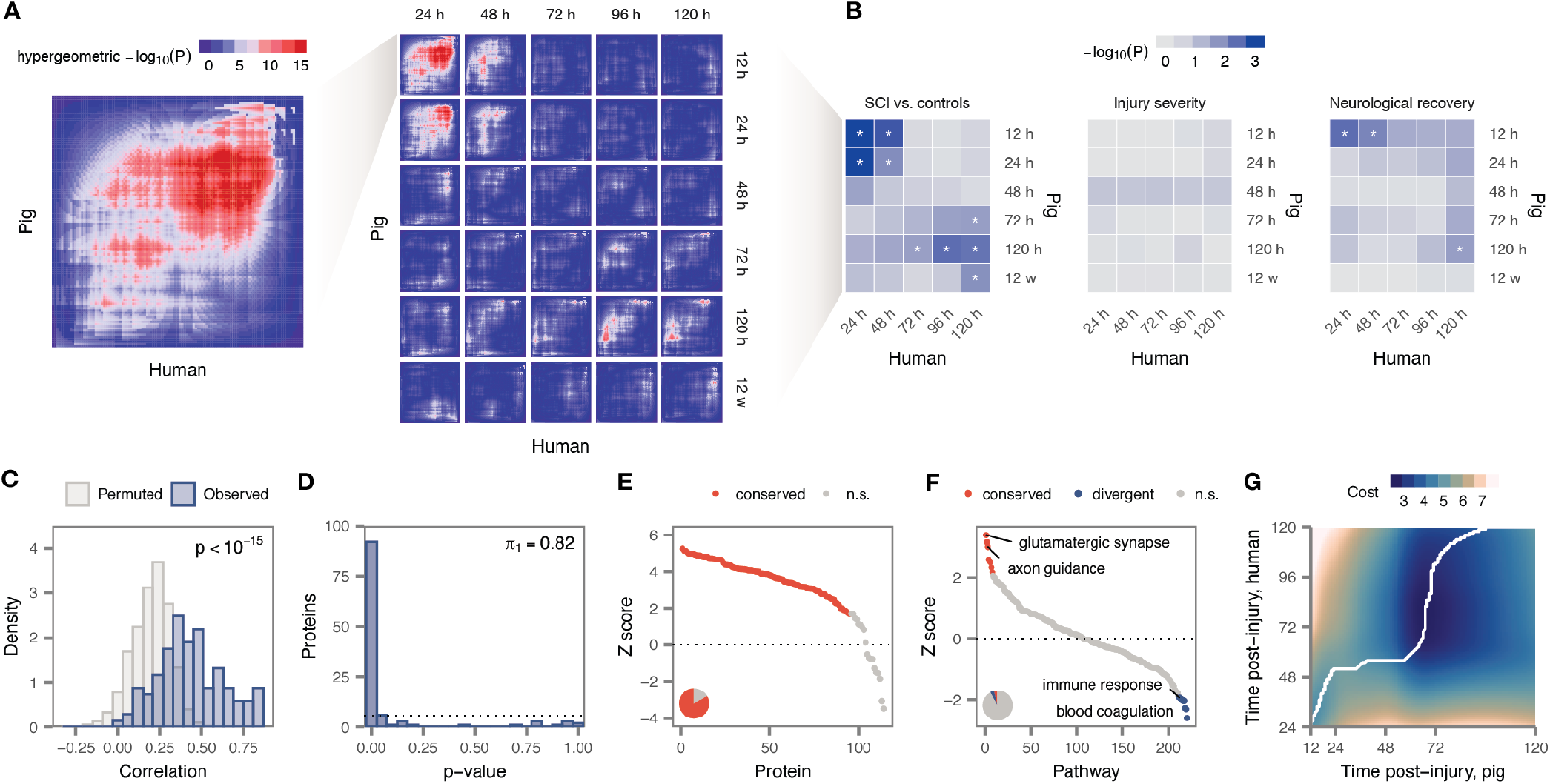
Evolutionary conservation of the response to SCI between humans and pigs. (A) Overview of the RRHO approach, comparing human and pig differential protein abundance after acute SCI. Left, RRHO plot comparing ranked lists of differentially abundant proteins at 24 h (human) and 12 h (pig) as an illustrative example. Right, RRHO plots for all pairs of timepoints in human and pig. (B) RRHO comparisons of differential protein abundance between human and pigs after acute SCI, left; as a function of baseline injury severity, middle; and as a function of neurological recovery, right. Asterisks denote comparisons with *p* < 0.05. (C) Distribution of NACC scores for proteins in human and pig samples, compared to randomly selected neighbor proteins. (D) Distribution of NACC score permutation p-values for individual proteins between human and pig. (E) Ranked NACC score z-scores for individual proteins, with significantly conserved proteins highlighted. Inset pie chart shows the total proportion of significantly conserved proteins. (F) Ranked NACC score z-scores for Gene Ontology functional categories, with significantly conserved pathways highlighted. Inset pie chart shows the total proportion of significantly conserved and divergent pathways. (G) Local cost matrix for the temporal alignment of human and pig CSF proteomes over the first five days post-injury by dynamic time warping; white line shows the optimal alignment.

We next applied RRHO to compare protein signatures of injury severity and neurological recovery (**Fig. 7B** and **Fig. S10A**). Broadly similar patterns to those identified for acute SCI were observed in the latter analysis, with the strongest overlap at the most acute stages of injury, and a more modest overlap at 120 h. In contrast, no significant overlaps were detected between human and pig with respect to baseline injury severity (**Fig. S10B**). This finding may reflect a fundamental dissimilarity between the precisely controlled measures of injury severity employed in animal studies, and the subjective, ordinal assessments carried out in the emergency room.

### Conserved and divergent patterns of protein co-expression

Beyond comparing signatures of specific clinical outcomes, we sought to identify individual proteins or biological processes with diverging patterns of abundance across species more generally. To this end, we applied neighborhood analysis of conserved coexpression (NACC) (*27*). This method has the advantage of not requiring that samples from either species be precisely matched with respect to clinical outcomes or timepoints, so long as two large and diverse panels are provided.

To apply NACC, we constructed protein co-expression networks in the CSF of either species, and evaluated the degree to which proteins tended to be co-expressed with the same neighbors across species. On a global level, we observed a significantly higher degree of shared co-expression between species than random expectation (*p <* 10^−15^, Brunner–Munzel test; **Fig. 7C**), providing further evidence for broad conservation of protein function in SCI. Using the *π*_1_ statistic, we estimated that 82% of proteins are co-expressed with shared neighbors across species (**Fig. 7D** and **Fig. S10C**). Ranking individual proteins based on the strength of conservation of their co-expression neighborhood (**Fig. 7E** and **Table S8A**), we found that no individual protein exhibited a significant divergence across species.

We then performed a similar NACC analysis at the level of biological processes. Among 220 GO terms robustly quantified in both species, eight were significantly conserved, whereas nine were divergent (**Fig. 7F** and **Table S8B**). GO terms displaying significantly conserved co-expression were largely related to neuronal structure and function (e.g., “axon guidance”, “glutamatergic synapse”). Conversely, GO terms associated with evolutionary divergence implicated immune and inflammatory processes (“immune response,” “cellular response to tumor necrosis factor”). Taken together, these analyses indicate broad conservation of protein function, but evolutionary divergence in the immune response to SCI.

### Temporal deviations in the SCI response across species

Preclinical studies in animal models often proceed from the implicit assumption that administering a therapeutic intervention at a particular timepoint in an animal model will elicit comparable effects at the same timepoint in humans. To directly address the possibility that comparable biological processes are activated at different timepoints in animal models, we applied dynamic time warping to temporally align the proteomes of both species over the first five days post-injury (**Fig. 7G**). Intriguingly, we found the two time courses were not directly aligned. Early timepoints in pig aligned to slightly later timepoints in human, suggesting that key aspects of the response to acute SCI progress faster in pig. This finding is consistent with RRHO analysis (**Fig. 7B**), where the strongest overlaps were observed between 12 h in pig and 24 h in human, and was further supported by inspecting alignments of individual proteins (**Fig. S10D**). For instance, the concentration of CKB in pig CSF decreases to a low at approximately 48 h, before returning to higher concentrations by 120 h (**Fig. S10E**); in humans, CKB had not yet returned to its initial concentration by 120 h. These observations provide systematic molecular evidence that the dynamics of the SCI response are not directly aligned between human patients and a large animal model, with particular implications for the neuroprotective agents that might be tested during the most acute phase of injury.

## Discussion

Despite the emergence of several promising therapeutic approaches in preclinical studies, clinicians presently lack effective treatments to restore function in acute SCI. In this work, we sought to address a number of factors underlying the translational divide in SCI through proteomic analyses of CSF and serum from 111 acute SCI patients and, in parallel, from a large animal model. Through our analysis of this dataset, we aimed to (i) enhance our basic understanding of the pathophysiology of acute human SCI, (ii) develop diagnostic and prognostic biomarkers of neurological impairment, and (iii) establish biological commonalities and differences between human SCI and a large animal model. Our findings have a number of implications for basic, translational, and clinical research in SCI.

Over the course of several decades, small-scale studies of SCI in animal models, primarily rodents, have amassed a substantial body of understanding about the pathobiology of SCI. In contrast, direct observations of secondary injury responses in human patients have historically been rare, and limited both by sample size and the number of analytes measured (*28*–*32*). The paucity of molecular-level descriptions of human SCI presents a broad limitation for translational research in a field that relies almost entirely on rodent experimental models for the development of new therapies. In this context, our analysis of 491 proteins across a total of 844 samples from 111 acute SCI patients provides an unprecedented resource to understand the evolving molecular responses to human SCI over the first five days post-injury, both within the central nervous system and systemically. Moreover, through serial collection of both biofluids over the first five days post-injury, we characterized the temporal profile of molecular changes following acute SCI, allowing us to dissect the complex biological responses that unfold over time after traumatic injury. Strikingly, we found that the proteomic alterations during the most acute phase, up to 24 h post-injury, were substantially distinct from those over the subsequent four days. This observation is particularly noteworthy given that our cross-species analysis established these early responses as most strongly conserved across species, and thus reveals an evolutionarily conserved ‘hyper-acute’ phase of SCI.

A primary goal of our study was to establish diagnostic and prognostic biomarkers of SCI. Through univariate analyses, we identified dozens of CSF and serum proteins with recurrent associations to injury severity or recovery, among which GFAP emerged as a particularly promising candidate. We then applied a machine-learning pipeline to develop multi-protein models. Perhaps most remarkably, we established a prognostic model capable of predicting AIS grade improvement at six months from the CSF proteome at 48 h post-injury with an AUC of 0.91. Excellent performance was also achieved in stratifying patients by AIS grade, identifying AIS A patients, and predicting motor complete injuries at six months. We subsequently went on to validate both univariate markers and preregistered multivariate models in an independent clinical cohort, in a manner designed to rigorously emulate a prospective clinical evaluation. Previous attempts to identify biomarkers in acute SCI have generally taken place in small patient cohorts, lacked validation in an independent cohort, and in several cases conflated training and test data when reporting accuracy. The methodological rigour of our efforts to validate these markers thus goes substantially beyond what has previously been achieved in this field. The success of this effort indicates that these tools could both facilitate the conduct of clinical trials, and inform decisions about acute and rehabilitative management, setting the stage for eventual precision medicine approaches.

The degree to which widely used animal models accurately reflect the pathophysiological processes active in SCI, and human disease more generally, has been extensively debated (*33*–*36*). Through comparative proteomic analysis in a porcine model, we provide systematic molecular evidence of substantial concordance in the molecular response to SCI between species, with several orthogonal analyses converging on an estimate that ~80% of proteome alterations are shared between human and pig. However, we also detected divergence in specific processes, most notably the immune response. We identified convergent proteomic signatures of neurological recovery, but not injury severity, possibly reflecting a fundamental dissimilarity between measures of severity employed in human and animal studies. Our search for conserved biological endpoints nonetheless led to the identification of GFAP as significantly correlated to both baseline severity and neurological outcome in both species. Last, our dense timecourse design in either species allowed us to perform, to our knowledge, the first systematic comparison of temporal patterns in the molecular response to a disease between species. Through a direct alignment of the human and pig time courses, we identified accelerated progression of the SCI response in our pig model, with implications for the timing of drug administration or other therapeutic interventions in preclinical studies. Collectively, this body of work addresses a number of outstanding questions about the validity of animal models of SCI, and defines a biochemical outcome measure for animal studies with direct relevance to the clinical setting.

Our study employed a targeted proteomics approach to measure the abundance of 491 proteins in the CSF and serum. The definitive advantage of this approach is its ability to provide precise quantification in analytically challenging biofluids, with sensitivity comparable to antibody-based assays (*37, 38*). Moreover, unlike these assays, this approach can be readily applied to proteins for which reliable antibodies may not exist. Conversely, although our interrogation of several hundred proteins within the CSF and serum of human SCI patients far exceeds any effort reported in the literature to date, our inferences are necessarily limited to the panel of proteins targeted here. An additional limitation of our study is the requirement that subjects complete a valid neurological examination within 24 h of admission; the possibility of important pathophysiological differences in patients unable to perform this exam cannot be excluded.

## Methods

### Clinical trial enrollment

Individuals sustaining an acute SCI were enrolled into this prospective observational study (clinicaltrials.gov: NCT01279811) at four North American sites if they met the following inclusion criteria: (1) AIS grade A, B, or C on presentation; (2) spinal injury between C1 and L1; and (3) the ability to provide a valid, reliable neurological examination within 24 h of injury. Patients were excluded if they had concomitant brain injuries or concomitant major trauma to the chest, pelvis, or extremities that required invasive intervention (e.g. chest tube, internal or external fixation), or were too sedated or intoxicated to provide a valid neurological examination. The study population was divided into two groups of consecutively enrolled patients, with the first 91 constituting the discovery cohort, and the subsequent 20 held out as a validation cohort. The uninjured control group consisted of 21 adult subjects undergoing routine lumbar decompressions and/or fusions who had no current or past history of SCI or myelopathy. The study was performed under the approval of the UBC Clinical Research Ethics Board (CREB; #H10-01091 for the clinical trial, #H08-00118 for normal controls).

Neurological evaluation. The severity of neurological impairment was graded according to the standards of the ISNCSCI examination, with motor scores recorded separately in the upper and lower extremities. All baseline testing and the assigning of the baseline AIS grade (A, B, or C) was conducted by research study nurses to confirm the initial examination of the patients. The baseline American Spinal Injury Association (ASIA) Impairment Scale (AIS) grade was A for 73 patients, B for 19, and C for a further 19, where AIS A denotes complete motor and sensory paralysis, AIS B denotes complete motor paralysis but some preserved sensation, and AIS C is assigned when there is some preserved motor and sensory function. The ISNCSCI examination was conducted again at six months postinjury in all but one patient, who was lost to follow-up and excluded from analyses of neurological recovery, to determine whether the AIS grade had improved and the extent of total motor score improvement.

### Porcine model

Female Yucatan miniature pigs (Sus scrofa) (Sinclair Bio-resources, Columbia, MO), aged approximately 150-200 days and weighing 20-30 kg, were group-housed at a large animal facility for 5 weeks prior to surgery. Animals were block-randomized into different injury severity groups, or a sham group. Animals in the sham group received an identical laminectomy surgery as in the injured groups, but no weight drop injury or compression to the spinal cord. Three different injury severities were induced by dropping a 50 g weight onto the exposed spinal cord from a height of 40, 20, or 10 cm, followed by 5 minutes of compression with a 150 g weight. Surgical procedures for experimental SCI and post-operative care were as previously described (*13, 39*). Briefly, using anatomical landmarks, the T9, T10, and T11 pedicles were cannulated and instrumented with screws (SelectTM Multi Axial Screw, Medtronic, Minneapolis, MN). After the T10 laminectomy was performed, the weight drop device was rigidly secured to the pedicle screws and positioned so that the impactor would fall directly on the exposed dura and spinal cord at T10. All animal experiments were conducted in accordance with the University of British Columbia Animal Care Committee (#A16-0311) and in strict adherence to the guidelines issued by the Canadian Council for Animal Care.

### Porcine neurological evaluation

The Porcine Thoracic Injury Behavior Scale (PTIBS) was used to assess hindlimb functional recovery, as previously described (*13*). Briefly, four weeks prior to injury, animals were trained to walk straight at a constant speed without stopping. Baseline behavior was obtained for each animal, one week prior to surgery; five runs were recorded using three high-definition camcorders placed 30 cm above the ground and behind the animals. Functional assessment resumed one week post-injury and continued once weekly for 12 weeks. The functional assessment footage was analyzed by two independent observers that were blinded to the biomechanical severity of spinal cord injury that was induced at the time of surgery. The PTIBS scale ranges from no active hindlimb movements (score 1), to normal ambulation (score 10). PTIBS scores of 1-3 are characterized by “hindlimb dragging,” scores of 4-6 reflect varying degrees of “stepping” ability, and scores of 7-10 reflect varying degrees of “walking” ability.

### Porcine histological evaluation

Tissue sparing was assessed by histological analysis at the end of the experiment (12 weeks post-injury), at which point animals were euthanized by sodium pentobarbital (120 mg/kg), the spinal cord harvested, post-fixed and cryoprotected as previously described (*40*). Subsequently, spinal cords were cut into 1 cm blocks centered on the site of injury, frozen on dry ice, and stored at −80°C. Cross-sections (20 *μ*m thick) were then cut using a cryostat. Sections were serially mounted onto adjacent silane-coated SuperFrost Plus slides (Fisher Scientific, Pittsburgh, PA) and stored at –80°C. For differentiating grey and white matter, Eriochrome Cyanine R staining (EC) was performed with Neutral Red as a counterstain. EC-stained sections were examined and micrographs (5 objective, Zeiss AxioImager M2 microscope) were taken of sections at 800 *μ*m intervals throughout the lesion site. The spinal cord outer perimeter, white matter, and gray matter were outlined, and the area of each was calculated using Zen Imaging Software (Carl Zeiss Canada Ltd., Toronto, ON, Canada). The percentages of white matter and grey matter were calculated by dividing the spared white or grey matter by the total area of the spinal cord on a given section, with the sum of the two representing spared tissue.

### Sample collection

Collection of CSF samples from enrolled patients was achieved using an intrathecal catheter (Braun Medical Inc., PA), inserted before the surgical procedure in the lumbar spine at L2/3 or L3/4 using a standard aseptic technique. The catheter was advanced 15-20 cm from the entry point on the skin surface and kept in place for five days. Samples of 3-4 mL were drawn at the time of catheter insertion and then in the subsequent postoperative period, approximately three times per day, discarding the first 1 mL of CSF aspirated. For the non-SCI control group, samples were acquired during the operation after the lumbar spine had been exposed. Sample processing was performed immediately at the patient’s bedside by the research study nursing team. CSF samples were centrifuged at 1,000 g for 10 min. Blood samples were incubated at room temperature for 25 min, then centrifuged at 10,000 g for 5 min. CSF and serum supernatants were collected and dispersed into 200 *μ*L aliquots, then immediately frozen in an ethanol-dry ice bath and stored at −80°C.

Serial collection of porcine samples was performed 15 minutes prior to injury, and then again at 12 h, 24 h, 48 h, 72 h, 120 h, and 12 w post-injury, as previously described (*15, 39*), and depicted in **Fig. S1**. Briefly, CSF collection was achieved using a 19 gauge epidural catheter (Braun Medical Inc., PA) inserted into the intrathecal space with the catheter tip resting approximately 8 cm caudal to the injury site. A total of 1 mL of CSF was collected and immediately centrifuged at 1,000 g for 10 minutes at room temperature. Serum collection was performed by inserting an 8F Groshong catheter (Bard Access Systems) in the left external jugular vein. This was connected to a low volume titanium subcutaneous access port (X-port; Bard Access Systems) housed in the posterior neck region. A total of 5 mL of whole blood was collected. To separate the serum portion, blood was allowed to incubate for 25 minutes at room temperature, and then centrifuged at 10,000 g for 5 minutes.

### Targeted proteomic assays

Serum protein quantitation was achieved using an expanded and updated version of a previously described multiple reaction monitoring (MRM) panel (*16, 17*) targeting 270 proteins. These peptides had been previously validated for their use in LC-MRM experiments following the CPTAC guidelines for assay development (*41*). Tryptic peptides were selected to serve as molecular surrogates for the 270 target proteins according to a series of peptide selection rules (e.g., sequence unique, devoid of oxidizable residues) and detectability, as described (*17*). All peptides were synthesized via Fmoc chemistry, purified through RP-HPLC with subsequent assessment by MALDI-TOF-MS, and characterized via amino acid analysis (AAA) and capillary zone electrophoresis (CZE).

For CSF protein quantitation, new parallel reaction monitoring (PRM) assays targeting 325 proteins were developed for the present study. CSF protein targets were selected on the basis of (1) an extensive literature review to identify proteins previously been implicated in SCI or traumatic brain injury (TBI), and (2) discovery proteomics experiments performed on undepleted CSF from injured patients and uninjured controls. Peptide targets from these proteins were selected using PeptidePicker (*42*) on the basis of detectability by mass spectrometry, digestion efficiency, lack of potential modifications, lack of potential interferences, and conservation between human and pig. A total of 500 isotopically labeled peptides, containing either ^13^C_6_^15^N_4_-Arg or ^13^C_6_^15^N_2_-Lys on the C-terminus of the peptide, were synthesized as above. These peptides were then individually validated for synthetic purity, detectability by mass spectrometry, and chromatographic performance. A total of 334 peptides were validated in this manner and were used for the remainder of the study. Collision energies and accumulation times were adjusted for individual peptides to provide optimal fragmentation (*43*) and adequate signal-to-noise ratios. High-quality tandem mass spectra were collected for each of these peptides, and a spectral library was produced containing all targets using Skyline (*44*).

### Sample processing

For serum samples, 10 *μ*L of raw serum was subjected to 9 M urea, 20 mM dithiothreitol, and 0.5 M iodoacetamide, all in Tris buffer at pH 8.0. Denaturation and reduction occurred simultaneously at 37°C for 30 min, with alkylation occurring thereafter in the dark at room temperature for 30 min. Proteolysis was initiated by the addition of TPCK-treated trypsin (35 *μ*L at 1 mg/mL; Worthington) at a 20:1 substrate:enzyme ratio. After overnight incubation at 37°C, proteolysis was quenched with 1% formic acid (FA). The SIS peptide mixture was then spiked into the digested samples, the standard curve samples, and the QC samples and concentrated by solid phase extraction (SPE) (Oasis HLB, 2 mg sorbent; Waters). After SPE, the concentrated eluate was frozen, lyophilized to dryness, and rehydrated in 0.1% FA (final concentration: 0.5 *μ*g/*μ*L digest) for LC-MRM/MS. For CSF samples, protein concentration was measured via Bradford assay, and one aliquot from each sample containing 200 *μ*g of protein was prepared. Samples were lyophilized and resuspended in 25 mM ammonium bicarbonate (ABC), and protein concentrations were validated by a second Bradford assay. Proteins were digested with trypsin as described (*45*). A mix of all SIS peptide standards was added, and the sample was cleaned on C18 desalting columns (MacroSpin, Nest Group, Southborough, MA). After C18 cleanup and drying, samples were resuspended in 60ul of 0.1% FA.

### Mass spectrometry

For serum samples, 20 *μ*L injections were separated with a Zorbax Eclipse Plus RP-UHPLC column (2.1 × 150 mm, 1.8 *μ*m particle diameter; Agilent) contained within a 1290 Infinity system (Agilent). Peptide separations were achieved at 0.4 mL/min over a 60 min run, via a multi-step LC gradient (2-80% mobile phase B; mobile phase compositions: A was 0.1% FA in water while B was 0.1% FA in acetonitrile). The column was maintained at 40°C. A post-gradient equilibration of 4 min was performed after each sample. The LC system was interfaced to a triple quadrupole mass spectrometer (Agilent 6490) via a standard-flow ESI source, operated in the positive ion mode. Peptide-specific LC-MS acquisition parameters were employed for optimal peptide ionization/fragmentation and scheduled MRM. Peptide optimizations were empirically optimized previously by direct infusion of the purified SIS peptides. Targets (one transition per peptide) were monitored over 500 ms cycles and 1 min detection windows.

For CSF samples, two 30 *μ*L injections were performed serially, one with each of two PRM acquisition methods. The two methods had identical ionization, calibration, autosampler, and liquid chromatography conditions. Specifically, peptides were separated over a 70 min run, using LC gradients as above; the separation column was a 2.1 × 250 mm C18 Poroshell column (Agilent), held at 45°; the flow rate was 100 *μ*L/min; and the ion source was the Agilent Jetstream ion source with reference mass calibration. The two methods differed in the peptide targets analyzed, with targets separated into the different methods based on retention time so as to minimize the number of concurrent eluting peptides analyzed per minute. Analysis of the CSF samples was performed on an Agilent 6550 QTOF mass spectrometer, coupled to an Agilent 1260 capillary flow HPLC, and an Agilent 1260 Autosampler.

### Data processing and quality control

Data from both serum and CSF samples was visualized and examined using Skyline (*44*). This involved peak inspection to ensure accurate selection, integration, and uniformity (in terms of peak shape and retention time) of the SIS and natural peptide forms. In the serum, a standard curve was prepared using a mix of light peptides that was spiked into a human tryptic digest in which the peptides were dimethylated (to shift their masses) from a high concentration of 1000 × the lower limit of quantitation (LLOQ) over 8 dilutions to the lowest point of the curve which was the LLOQ for the assay. The QC samples were prepared from the same light peptide mix and diluted in dimethylated human digest at 4×, 50×, and 500× the LLOQ for each peptide. After defining a small number of criteria (i.e., 1/x regression weighting, *<*20% deviation in the QC level accuracy) the standard curve was used to calculate the peptide concentration in fmol/*μ*l in the patient samples through linear regression. Protein quantitations below or above the LOQ were removed. In the CSF, peak ratios were calculated and extracted using the top six interference-free fragment ions that matched the library spectra. Protein quantitations were removed when the dot product to the library spectrum was less than 0.7 or the maximum peak height was below 20.

### Modularity analysis

Modularity analysis was performed essentially as previously described (*18*), using the R package ‘igraph’ (*46*). Given a network and a grouping of nodes, modularity measures the degree to which nodes within each group are preferentially connected to one another. A weighted network was constructed in which nodes represent samples, and an edge was drawn between pairs of nodes when the Pearson correlation coefficient, computed across all proteins quantified in both samples, was higher than a given threshold. The modularity was subsequently computed with respect to several potential groupings of nodes (including patient, level of injury [cervical, thoracic, or lumbar], injury severity [base-line AIS grade], sex, or time post-injury) as a function of network density, defined as the proportion of possible edges used to construct the network.

### Univariate analyses

Univariate analysis of protein abundance was performed using linear models with empirical Bayes shrinkage as implemented in the R package ‘limma’ (*47*), and p-values were adjusted using Benjamini-Hochberg correction to control the false discovery rate. The plate on which each sample was stored and processed was included in all models in order to guard against the possibility of batch effects. No imputation of missing values was performed for univariate analysis.

In the human dataset, associations between protein abundance and three clinical variables were evaluated: (i) injury status (i.e., comparing samples from patients with acute SCI to uninjured controls); (ii) the severity of the initial injury, assessed using the AIS scale; (iii) recovery of motor function at six months post-injury, quantified by the change in ASIA total motor score (ΔTMS); and improvement in AIS grade at six months. Baseline AIS grade was modeled as an ordinal variable, ΔTMS as a continuous variable, and AIS grade improvement as a binary variable. Baseline neurological injury level (e.g., C6 vs. T9) was included as a co-variate in models of six-month ΔTMS and AIS grade improvement, in order to account for inherently different absolute potentials for motor recovery.

In the pig dataset, associations between protein abundance and four experimental variables were evaluated: (i) injury status (i.e., comparing samples drawn from pigs after experimental SCI to matched samples from the same pigs at baseline); (ii) base-line injury severity, quantified by the height of the weight-drop contusion/compression impactor; (iii) hindlimb neurological recovery, quantified using the average Porcine Thoracic Injury Behavior Scale (PTIBS) score across weeks 10-12 post-injury; and (iv) white matter tissue sparing, quantified by histology at 12 weeks post-injury. Baseline injury severity was modeled as an ordinal variable, whereas hindlimb neurological recovery and white matter tissue sparing were modeled as continuous variables.

### Consensus clustering

*k*-means consensus clustering (*20*) was performed on the matrix of protein log_2_-fold change values over the first five days post-injury, using the R package ‘ConsensusClusterPlus’ (*48*). The Euclidean distance was used as the distance function, and 100 subsamples were performed. The change in the area under the cumulative distribution function (AUCDF) was used to identify the optimal number of clusters, as the value of *k* at which there was no appreciable increase in the AUCDF. This procedure yielded a total of nine clusters in the CSF and eight in the serum. Of these, two in the CSF and one in the serum comprised two or fewer proteins and were removed, resulting in seven clusters of temporally co-regulated proteins in each biofluid. Gene Ontology terms enriched within each cluster, relative to the remaining clusters, were identified using the conditional hypergeometric test (*49*) implemented in the R package ‘GOstats’ (*50*).

### Multivariate analysis

We implemented an extensive pipeline to develop multivariate diagnostic and prognostic models using the python package ‘scikit-learn’ (*51*), encompassing both parametric (e.g., penalized logistic regression) and non-parametric (e.g., gradient boosting machine) approaches. Specifically, f or b inary or multi-class classification tasks (including baseline AIS grade, base-line AIS A, change in TMS of seven or more points, AIS grade improvement, and motor complete injury at six months), we evaluated twelve different families of models, including Gaussian naive Bayes, *k*-nearest neighbors, nearest-centroid, support vector machine, logistic regression, linear discriminant analysis, random forest, extra trees, adaptive boosting, gradient boosting, and XGBoost classifiers. For the lone regression task, change in TMS at six months, we evaluated a further eleven families of models, including *k*-nearest neighbor, support vector machine, penalized logistic regression (with L1, L2, or elastic net penalties), L1-penalized least angle regression, random forest, extra trees, adaptive boosting, gradient boosting, and XGBoost regressors. Model performance was evaluated in five-fold cross-validation, using the accuracy and balanced accuracy to evaluate multi-class prediction, the AUC and accuracy to evaluate binary prediction, and the coefficient of determination (r^2^ score) to evaluate regression. Hyperparameter grids were adapted from Olson et al. (*52*), with minor modifications, and are provided in **Table S9**. In addition to evaluating model hyperparameters, we also evaluated preprocessing choices, including scaling and feature selection, using the scikit-learn ‘Pipeline’ class. Feature selection was optionally performed by using limma to identify peptides with significant univariate associations to the clinical outcome of interest at nominal significance (that is, uncorrected *p* < 0.05), and was pre-calculated for each cross-validation fold in order to prevent inadvertent leakage of information between training and test splits. We additionally evaluated the impact of providing clinical data from the baseline neurological examination in conjunction with proteomic data, including age, sex, neurological level of injury, baseline AIS grade, baseline UEMS, baseline LEMS, and baseline AMS. Categorical features from the baseline neurological examination (baseline AIS grade and level of injury) were converted to binary indicator variables.

For the analysis of the validation cohort, the best models as determined by cross-validation in the discovery cohort were re-trained on the entirety of the discovery cohort, and projected into the validation cohort to predict clinical outcomes. A total of six models were selected and pre-registered for validation on the basis of the discovery cohort analysis. Samples from the validation cohort were blinded to investigators until the finalization of all statistical analyses of the discovery cohort, in order to provide an independent, blinded validation set.

### Cross-species analysis

Unless otherwise noted, all cross-species analyses were performed on a combined matrix of human and pig protein abundance including all proteins quantified in at least one-third of samples from either species, which was normalized by peptide as described (*53*). Modularity analysis was performed as described above, after restricting samples to those collected at four timepoints overlapping between species (24 h, 48 h, 72 h, and 120 h). The proportion of true null hypotheses, *π*0, was estimated using the R package ‘qvalue’ (*25*), from which the estimated proportion of true associations, *π*_1_, was calculated as 1 − *π*_0_.

Rank-rank hypergeometric overlap analysis30 was performed using the R package ‘RRHO’. Briefly, after performing independent univariate analyses in either species as described above, all proteins quantified in either species were ranked by the strength and direction of DE, and a pair of sliding windows were passed along each of the two ranked lists. The hypergeometric probability of the observed overlap between proteins in each species was calculated for each pair of ranks. The statistical significance of each overlap was calculated by randomly permuting the status (e.g., SCI vs. control) of each human and pig sample, performing an identical univariate analysis with the permuted data, and comparing the maximum – log10 p-values achieved in 1,000 permutations to the observed value (*54*).

Neighborhood analysis of conserved coexpression (NACC) (*27*) was performed to estimate the overall conservation of protein abundance patterns in SCI between species, and to identify individual proteins or specific pathways with conserved or divergent patterns of abundance. Briefly, independent protein co-expression networks were constructed in either species, using the Pearson correlation coefficient to quantify similarity. The *k* nearest neighbors of each protein in turn were identified in the human co-expression network, and the mean correlation coefficient between the human neighbors and the protein of interest were calculated in pig. This procedure was repeated with human and pig inverted, and the average correlation across the two comparisons was retained as a symmetric measure of the degree of conservation of co-expression for each protein. We used a neighborhood size of *k* = 10, but found our conclusions robust to the neighborhood size (**Fig. S10C**). Permutation p-values (*54*) were calculated for each individual protein by comparing the observed NACC score to a distribution of NACC scores derived from 1,000 random sets of ‘neighbors’. An identical permutation test was applied to GO terms annotated to between three and 50 proteins in both species, using the average NACC score of all genes within each GO category as the test statistic as previously described (*27*).

Dynamic time warping was performed using the R package ‘dtw’ (*55*). Samples from uninjured controls and porcine samples collected at 12 weeks post-injury were removed. Abundance profiles for each protein in either species were smoothed using a local polynomial filter to enhance time resolution (*56*), after which pair-wise Euclidean distances were calculated between human and pig protein abundance at each pair of timepoints, and supplied as the local cost matrix to identify the optimal global alignment.

## Data availability

The raw mass spectrometry data have been deposited to the ProteomeXchange Consortium (*57*) via the MassIVE partner repository with the dataset identifier MSV000085567 (reviewer access token: porcinetohuman). Clinical and technical metadata and processed protein abundance data are provided in **Tables S1** and **S3**, respectively.

## Acknowledgements

This work was supported by Brain Canada, the Michael Smith Foundation for Health Research, Genome Canada, Genome BC, the Djavad Mowfaghian Center for Brain Health, the Praxis Spinal Cord Institute (formerly the Rick Hansen Institute), and the VGH & UBC Hospital Foundation. The analysis was enabled in part by the support provided by WestGrid and Compute Canada (to L.J.F.), and through computational resources and services provided by Advanced Research Computing at the University of British Columbia (to B.K.K. and L.J.F.). The authors are grateful for the expertise of the dedicated staff at the UBC Center for Comparative Medicine who helped to manage the health and welfare of the animals. M.A.S. acknowledges support from the Canadian Institutes of Health Research (CIHR), a Vancouver Coastal Health–CIHR–UBC MD/PhD Studentship, and the Wings for Life Spinal Cord Research Foundation. Mass spectrometry infrastructure for L.J.F. and C.B. was supported by Genome BC and Genome Canada (214PRO). B.K.K. is the Canada Research Chair in Spinal Cord Injury and the Dvorak Chair in Spine Trauma.

---

**Fig. S1:**
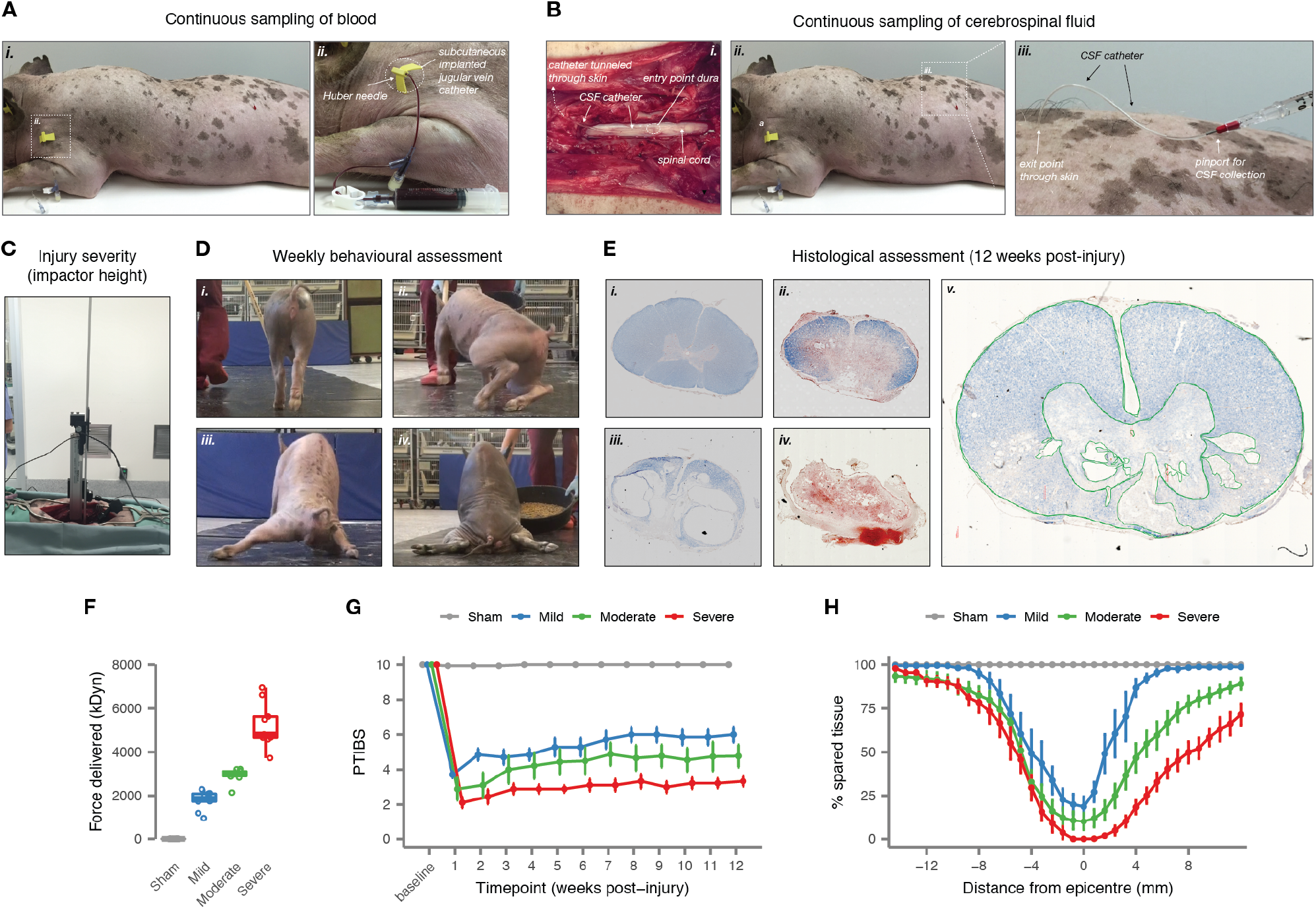
Experimental study of SCI in a large animal model, the Yucatan pig. (A-B) Overview of surgical and experimental setup for continuous collection of CSF (A) and serum (B) samples. (C-E) Overview of key experimental outcomes. (C) Baseline injury severity was quantified by impactor height (mild, 10 cm; moderate, 20 cm; severe, 40 cm). (D) Neurological recovery was quantified by weekly behavioral assessments using the Porcine Thoracic Injury Behavioral Scale (PTIBS) for hindlimb function. Representative images from PTIBS assessments are shown for sham (i), mild (ii), moderate (iii), and severe (iv) injuries. (E) Tissue sparing was quantified by histology at 12 weeks post-injury, on the basis of cross-sections of the spinal cord from 13.6 mm rostral to 13.6 mm caudal to the lesion site, in increments of 0.8 mm. Slides were manually traced to determine the extent of spared tissue (v). (F) Maximum force delivered, in kDyn, to animals in each of four treatment groups. (G) Mean PTIBS scores for animals in each of four treatment groups over the first twelve weeks post-injury. Bars show standard error. (H) Mean proportion of spared tissue in sections taken from 13.6 mm rostral to 13.6 mm caudal to the lesion, relative to the entire area of the spinal cord on histological section. Bars show standard error.

**Fig. S2:**
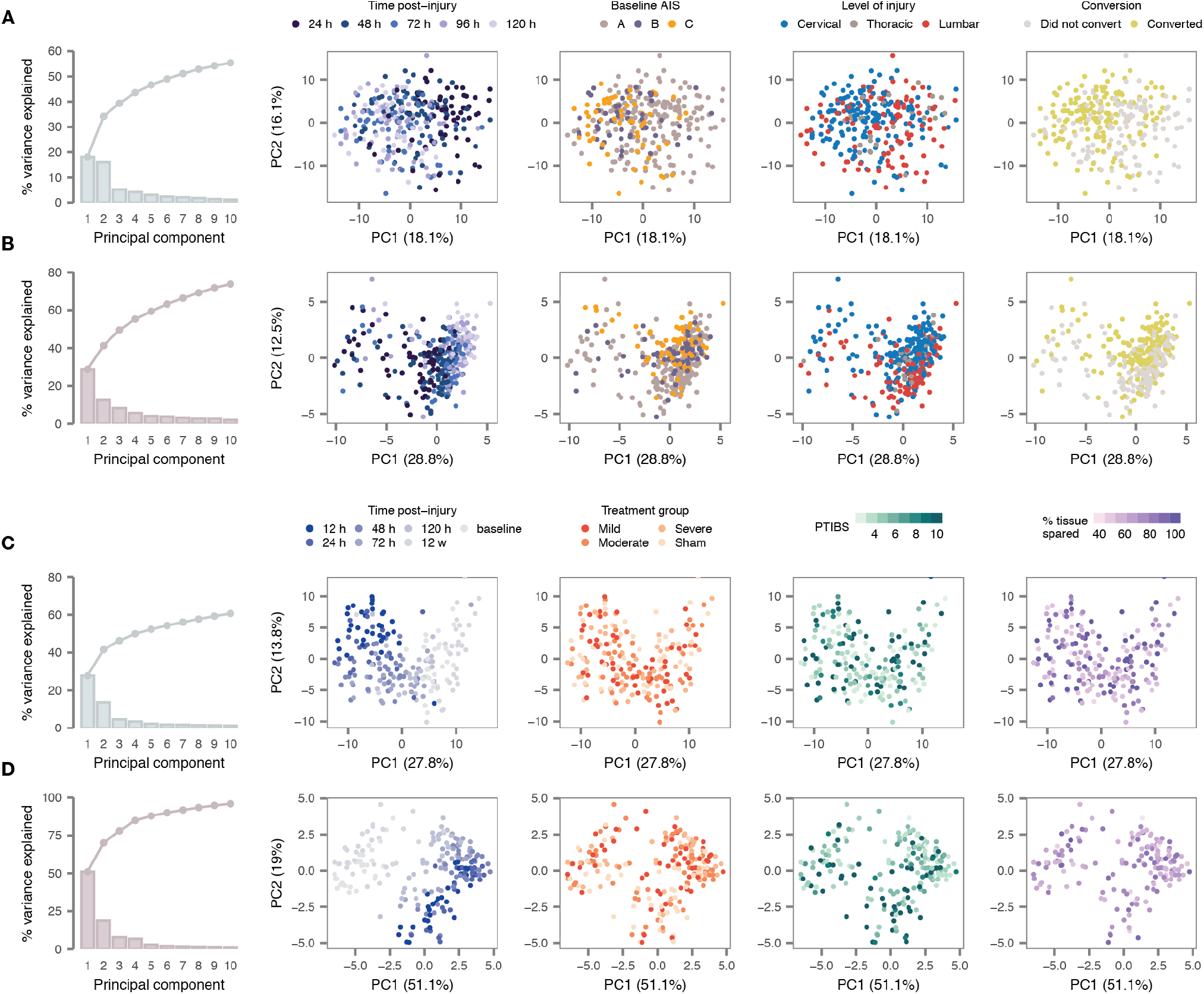
Principal component analysis of CSF and serum proteomes in human and pig. (A) Principal component analysis of CSF proteomics data from the human cohort. Left, percent of variance explained by principal components 1 to 10 (bars) or cumulatively by the first 1 to 10 principal components (line). Right, biplot of the first two principal components (PC1 and PC2) with samples coloured by time post-injury, baseline AIS grade, level of injury, or AIS conversion. (B) As in (A), but for the serum proteomics data from the human cohort. (C) Principal component analysis of CSF proteomics data from the pig cohort. Left, percent of variance explained by principal components 1 to 10 (bars) or cumulatively by the first 1 to 10 principal components (line). Right, biplot of the first two principal components (PC1 and PC2) with samples coloured by time post-injury, treatment group (baseline injury severity), PTIBS score at 12 weeks post-injury, and percentage of tissue spared as assessed by histology at 12 weeks post-injury. (D) As in (C), but for the serum proteomics data from the pig cohort.

**Fig. S3:**
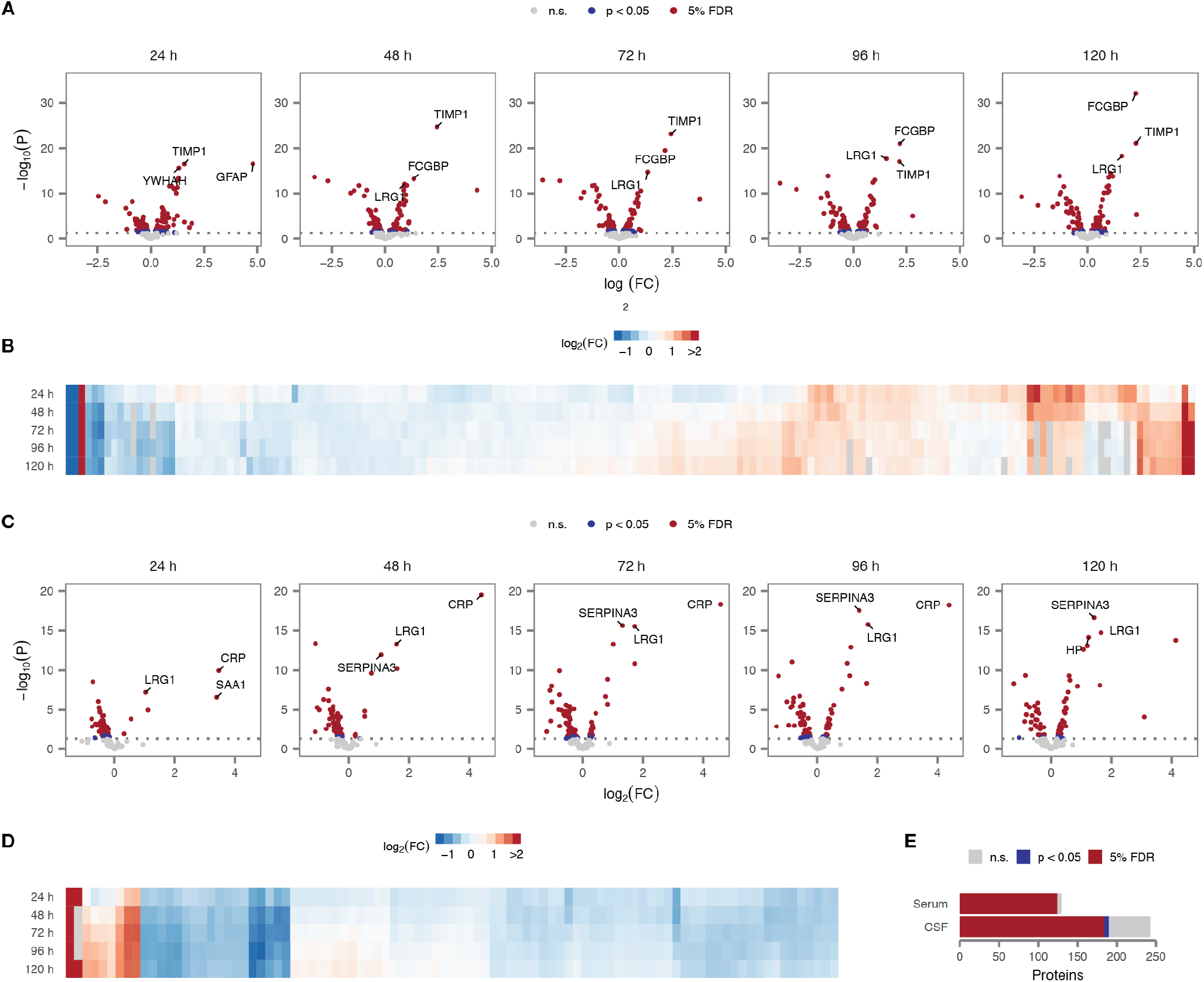
Alterations in CSF and serum protein abundance in acute SCI. (A) Volcano plots of differential protein abundance between 24 h and 120 h post-injury in CSF samples from patients with acute SCI compared to uninjured controls. (B) Time courses of differential protein abundance (log-fold change, relative to uninjured controls) over the first five days post-injury for 171 CSF proteins differentially expressed between SCI and control samples within at least one timepoint. (C) As in (A), but showing differential abundance of serum proteins compared to uninjured controls. (D) As in (B), but showing time courses of 92 serum proteins differentially expressed between SCI and control samples within at least one timepoint. (E) Numbers of CSF and serum proteins that display a statistically significant association with time post-injury.

**Fig. S4:**
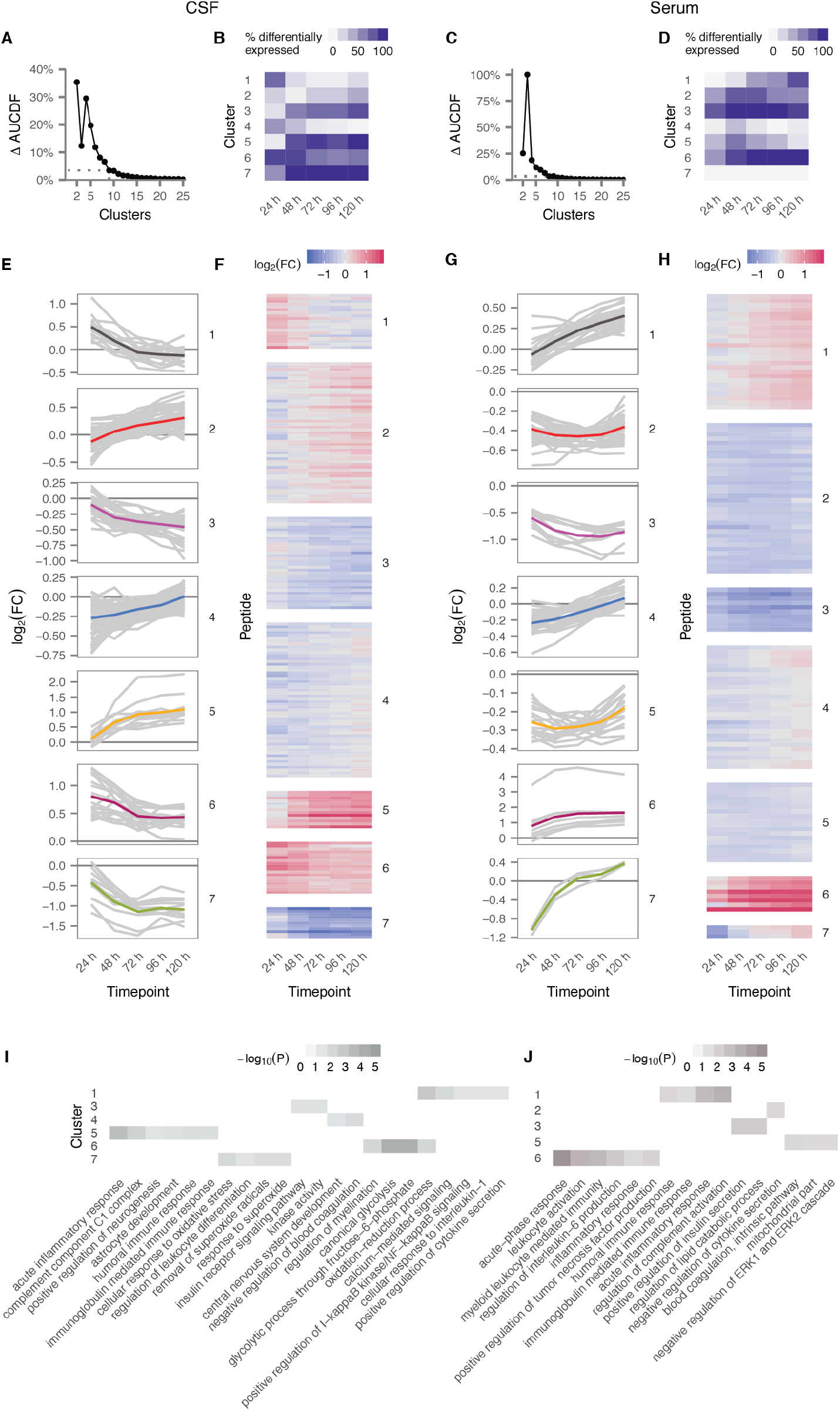
Modules of temporally co-regulated CSF and serum proteins in acute SCI. (A) Relative change in the area under the cumulative distribution function (*20*) comparing consensus clustering solutions of *k* and *k* –1 clusters of the CSF proteome. The optimal number of clusters was determined for each tissue as the value of *k* at which there was no appreciable increase in the AUCDF. (B) Proportion of peptides from each cluster found to be differentially expressed between acute SCI patients and uninjured controls at each timepoint. (C-D) As in (A-B), but for the serum proteome. (E) Time courses of differential protein abundance (log-fold change, relative to uninjured controls) over the first five days post-injury for peptides in each of the seven CSF protein modules, grey lines, and the mean time course for the entire module, colored line. (F) Time courses of differential protein abundance, as in **c,** shown as a heatmap. (G-H) As in (E-F), but for the serum proteome. (I-J) GO terms enriched in each of each of the seven CSF (I) and serum (J) protein modules.

**Fig. S5:**
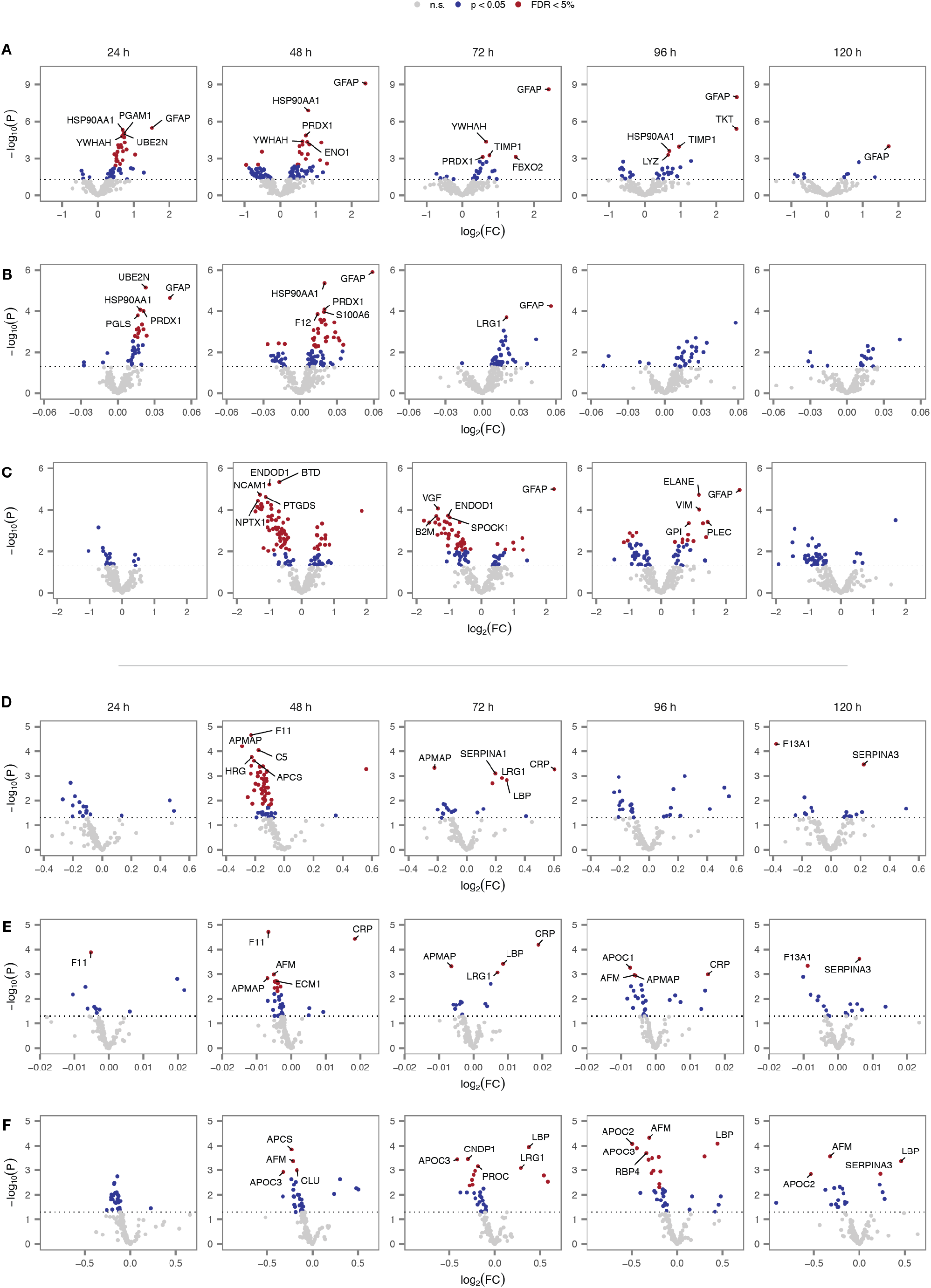
Univariate analysis of injury severity and neurological recovery in human serum and CSF. (A) Volcano plots of differential protein abundance as a function of injury severity, as quantified by the baseline AIS grade, in CSF samples between 24 h and 120 h post-injury. (B) Volcano plots of differential protein abundance as a function of neurological recovery at six months post-injury, as quantified by the change in total motor score relative to baseline, in CSF samples between 24 h and 120 h post-injury. (C) Volcano plots of differential protein abundance as a function of neurological recovery at six months post-injury, as quantified by improvement in the AIS grade relative to baseline, in CSF samples between 24 h and 120 h post-injury. (D) As in (A), but for serum samples. (E) As in (B), but for serum samples. (F) As in (C), but for serum samples.

**Fig. S6:**
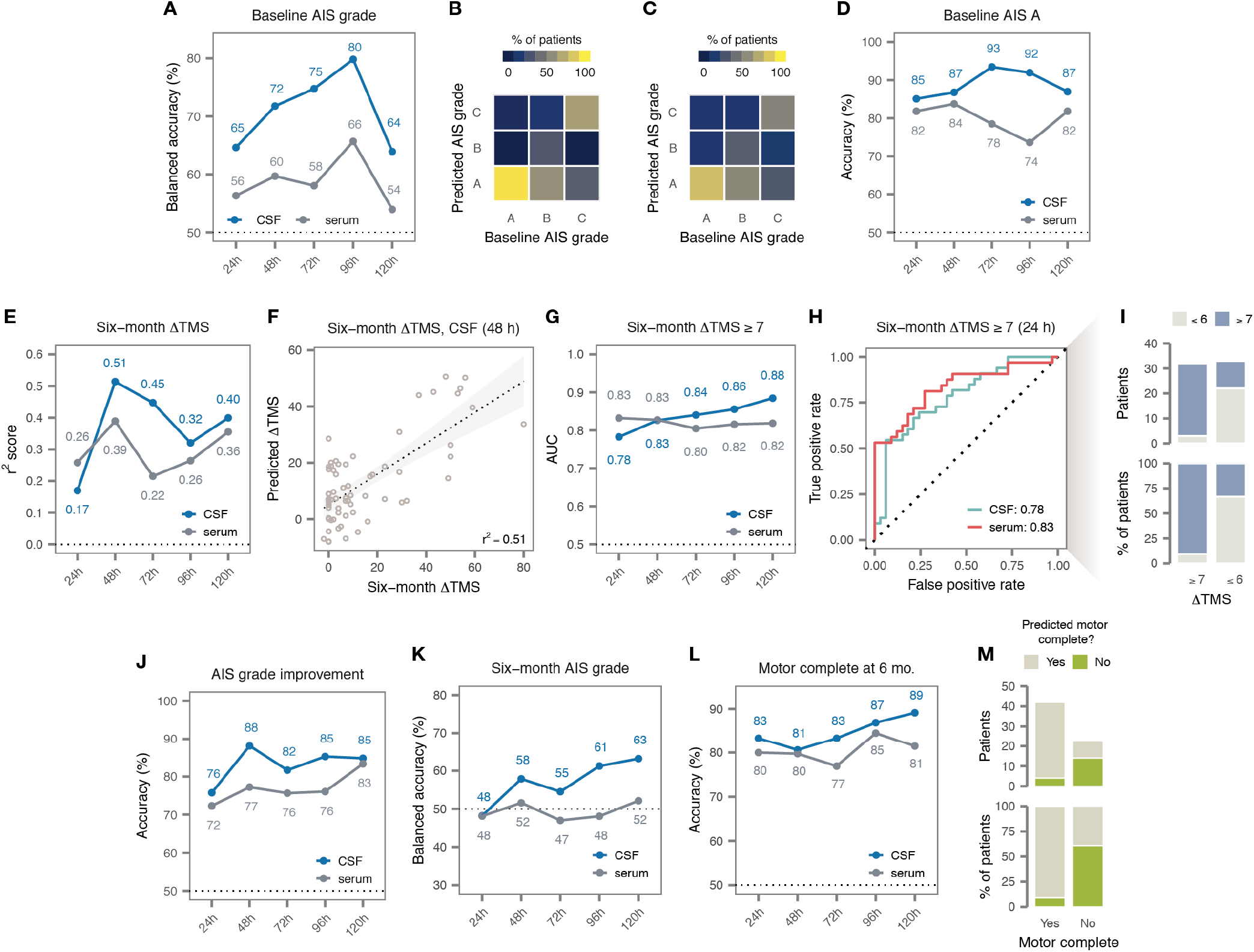
Additional multivariate analysis of SCI severity and recovery. (A) Cross-validation balanced accuracy of multivariate diagnostic models trained to stratify patients by baseline AIS grade, at timepoints between 24 and 120 h post-injury. (B) Confusion matrix of the best CSF diagnostic model of baseline AIS grade at 24 h post-injury. (C) Confusion matrix of the best serum diagnostic model of baseline AIS grade at 24 h post-injury. (D) Cross-validation accuracy of multivariate diagnostic models trained to discern patients with a baseline AIS grade of A, at timepoints between 24 and 120 h post-injury. (E) Cross-validation coefficient of determination (r^2^ score) of multivariate prognostic models trained to predict the change in TMS at six months post-injury, relative to baseline, at timepoints between 24 and 120 h post-injury. (F) Predictions made by the best CSF prognostic model of six-month change in TMS at 48 h post-injury. (G) Cross-validation AUC of multivariate prognostic models trained to predict a change in TMS of seven or more points at six months post-injury, at timepoints between 24 and 120 h post-injury. (H) ROC curves of the best CSF and serum prognostic models of six-month change in TMS of seven or more points at 24 h post-injury. (I) Predictions made by the best serum prognostic model at 24 h post-injury. (J) Cross-validation accuracy of multivariate prognostic models trained to predict improvement in AIS grade at six months post-injury, relative to baseline, at timepoints between 24 and 120 h post-injury. (K) Cross-validation balanced accuracy of multivariate prognostic models trained to predict AIS grade at six months, at timepoints between 24 and 120 h post-injury. (L) Cross-validation accuracy of multivariate prognostic models trained to predict motor complete vs. incomplete injury at six months, at timepoints between 24 and 120 h post-injury. (M) Predictions made by the best serum prognostic model of motor complete vs. incomplete injury at six months, at 24 h post-injury.

**Fig. S7:**
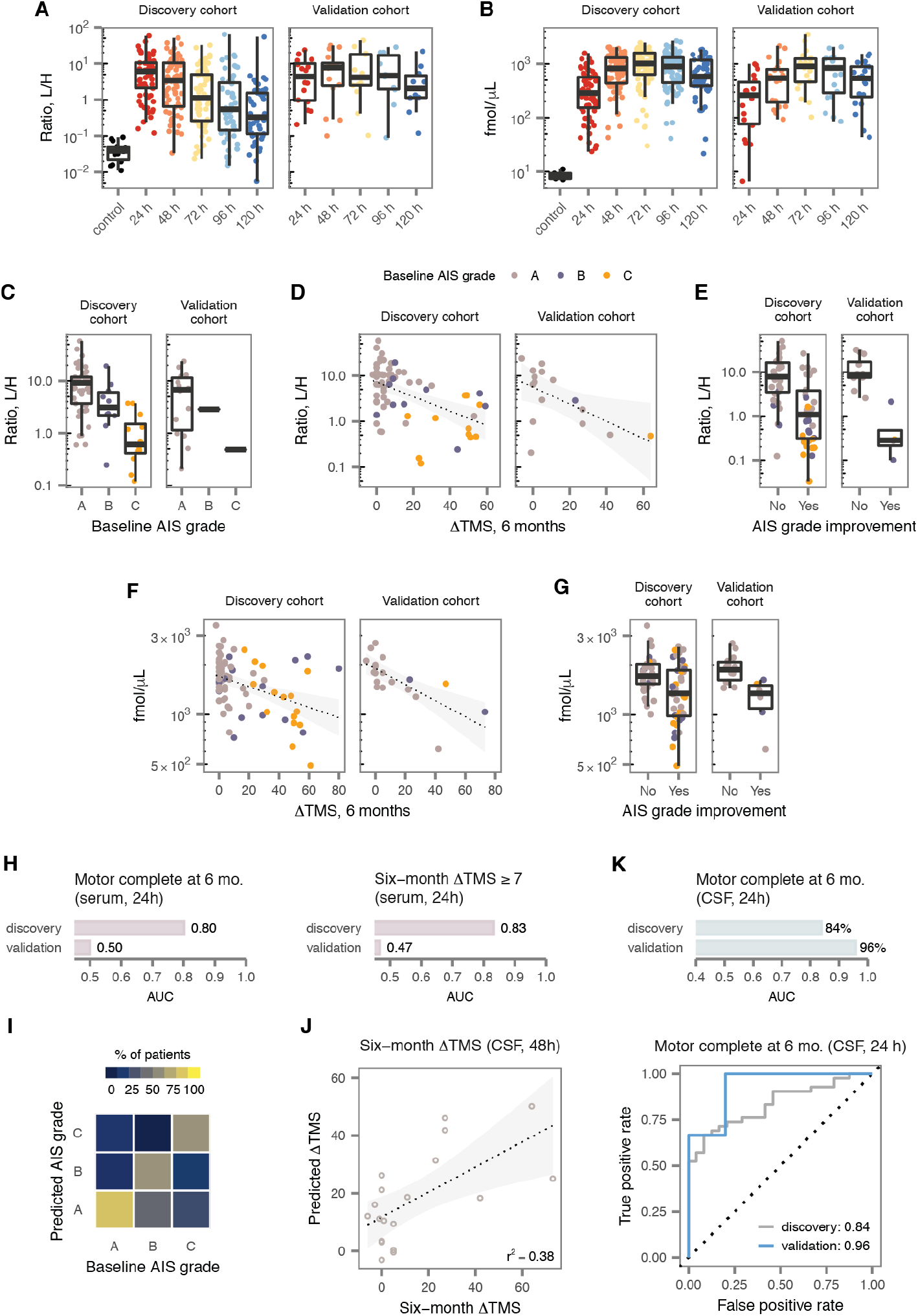
Replication of univariate associations and multivariate models in an independent validation cohort. (A) Time course of CSF abundance for an exemplary protein with replicated alterations between acute SCI patients and uninjured controls, GFAP. (B) Time course of serum abundance for an exemplary protein with replicated alterations between acute SCI patients and uninjured controls, CRP. (C-G) Examples of proteins with univariate associations to severity or recovery replicated in the validation cohort. (C) Abundance of GFAP in CSF samples at 24 h, stratified by baseline AIS grade. (D) Abundance of GFAP in CSF samples at 24 h, stratified by change in TMS at six months post-injury. (E) Abundance of GFAP in CSF samples at 48 h, stratified by improvement in AIS grade at six months post-injury. (F) Abundance of LRG1 in serum samples at 72 h, stratified by change in TMS at six months post-injury. (G) Abundance of LRG1 in serum samples at 72 h, stratified by improvement in AIS grade at six months post-injury. (H) Performance of two preregistered serum multivariate models in the validation cohort. (I) Confusion matrix of the best CSF diagnostic model of baseline AIS grade at 48 h post-injury in the validation cohort. (J) Performance of the best CSF prognostic model of six-month change in TMS at 48 h post-injury in the validation cohort. (K) Performance, top, and ROC curve, bottom, of the best CSF prognostic model of motor complete vs. incomplete injury at six months at 24 h post-injury.

**Fig. S8:**
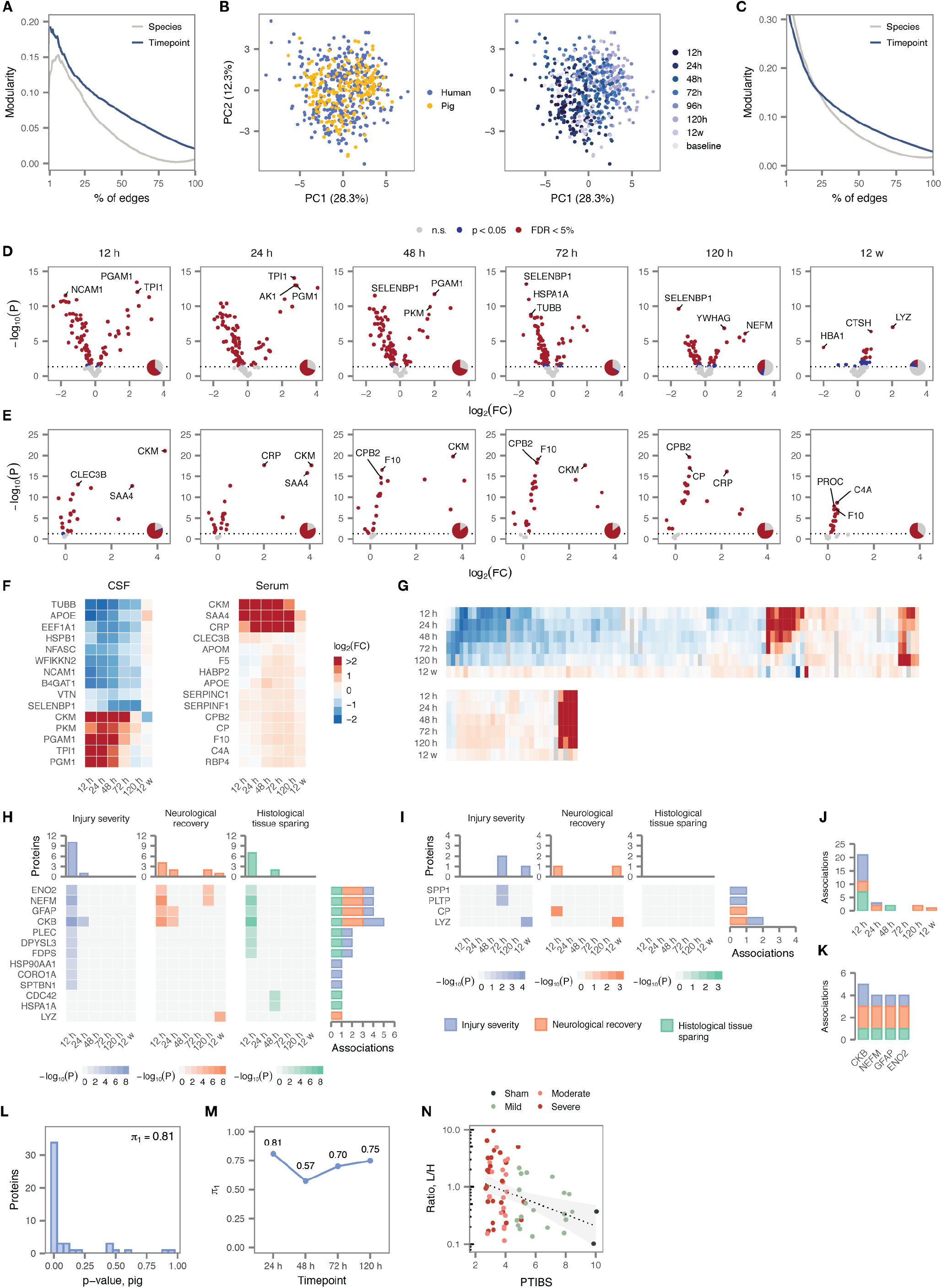
Univariate analysis of acute SCI, injury severity, and neurological recovery in pig serum and CSF. (A) Modularity analysis of the combined human and pig CSF proteomes, with samples grouped by species or time post-injury. Modularity is shown as a function of the number of edges between samples used to construct the network, as a percentage of the total number of possible edges. (B) Principal component analysis of human and pig serum proteomes, with samples colored by species, left, or time post-injury, right. (C) As in (A), but for the combined serum proteomes. (D) Volcano plots of differential protein abundance between 24 h and 120 h post-injury in CSF samples from injured pigs, compared to samples drawn from the same pigs at baseline (15 min prior to injury). (E) As in (D), but for serum protein abundance. (F) Time courses of differential protein abundance (log2-fold change, relative to baseline) over twelve weeks post-injury for fifteen of the most profoundly altered CSF proteins, left, and serum proteins, right. (G) Time courses of differential protein abundance (log2-fold change, relative to uninjured controls) over twelve weeks post-injury for 135 CSF proteins, top, and 26 serum proteins, bottom, differentially expressed between SCI and control samples within at least one timepoint. (H) Statistical significance of associations between CSF protein abundance and three experimental variables over twelve weeks post-injury for 13 proteins with at least one significant association. Grey squares indicate associations that were not significant after correction for multiple hypothesis testing. (I) As in (H), but for four serum proteins with at least one significant association. (J-K) Number of statistically significant associations between CSF protein abundance and three experimental variables outcomes per timepoint (J), and for the four CSF proteins with the most recurrent associations (K). (L) Distribution of p-values for proteins differentially expressed in the CSF of acute SCI patients vs. uninjured controls, in comparisons of pig CSF samples drawn at 24 h post-injury and at baseline. (M) Estimated proportion of true associations, *π*_1_, for all human-pig comparisons between 24 h and 120 h. (N) Abundance of GFAP in pig CSF samples at 24 h, stratified by PTIBS score.

**Fig. S9:**
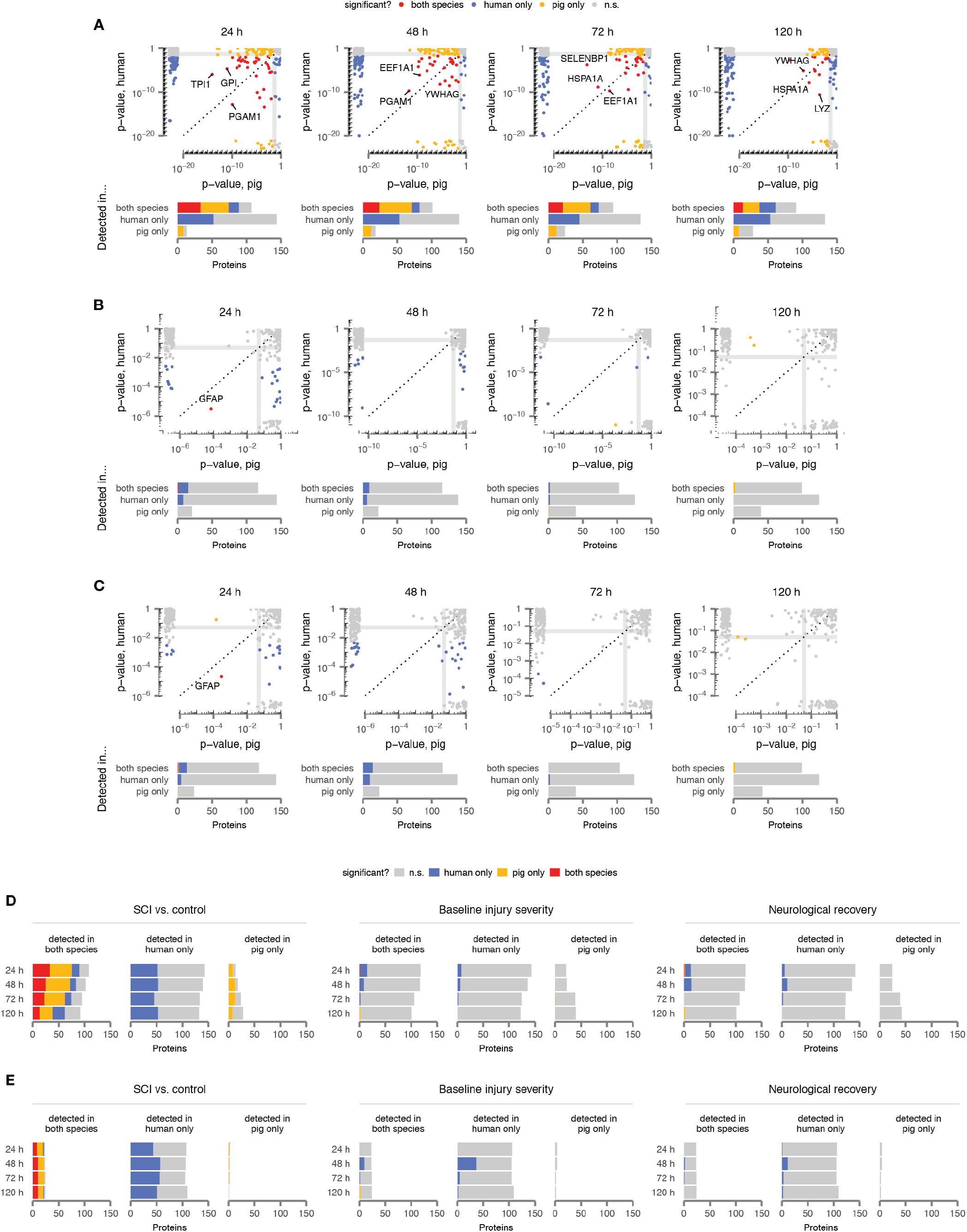
Overlap of differentially abundant proteins between human and pig. (A) Top, p-values for differential protein abundance between samples from individuals with acute SCI and uninjured controls at four matching timepoints in human and pig. Marginal plots show p-values for proteins quantified in human (pig) only. Bottom, number of proteins with statistically significant differential abundance in both species, human only, pig only, or neither, among proteins quantified in both species, human only, or pig only. (B) As in (A), but for proteins associated with injury severity. (C) As in (A), but for proteins associated with neurological recovery. (D) Summary of significant univariate associations involving CSF proteins across species, stratified by (i) clinical or experimental outcome, (ii) time post-injury, (iii) detection in one or both species, and (iv) statistical significance in neither, one, or both species. (E) As in (D), but for the serum proteome.

**Fig. S10:**
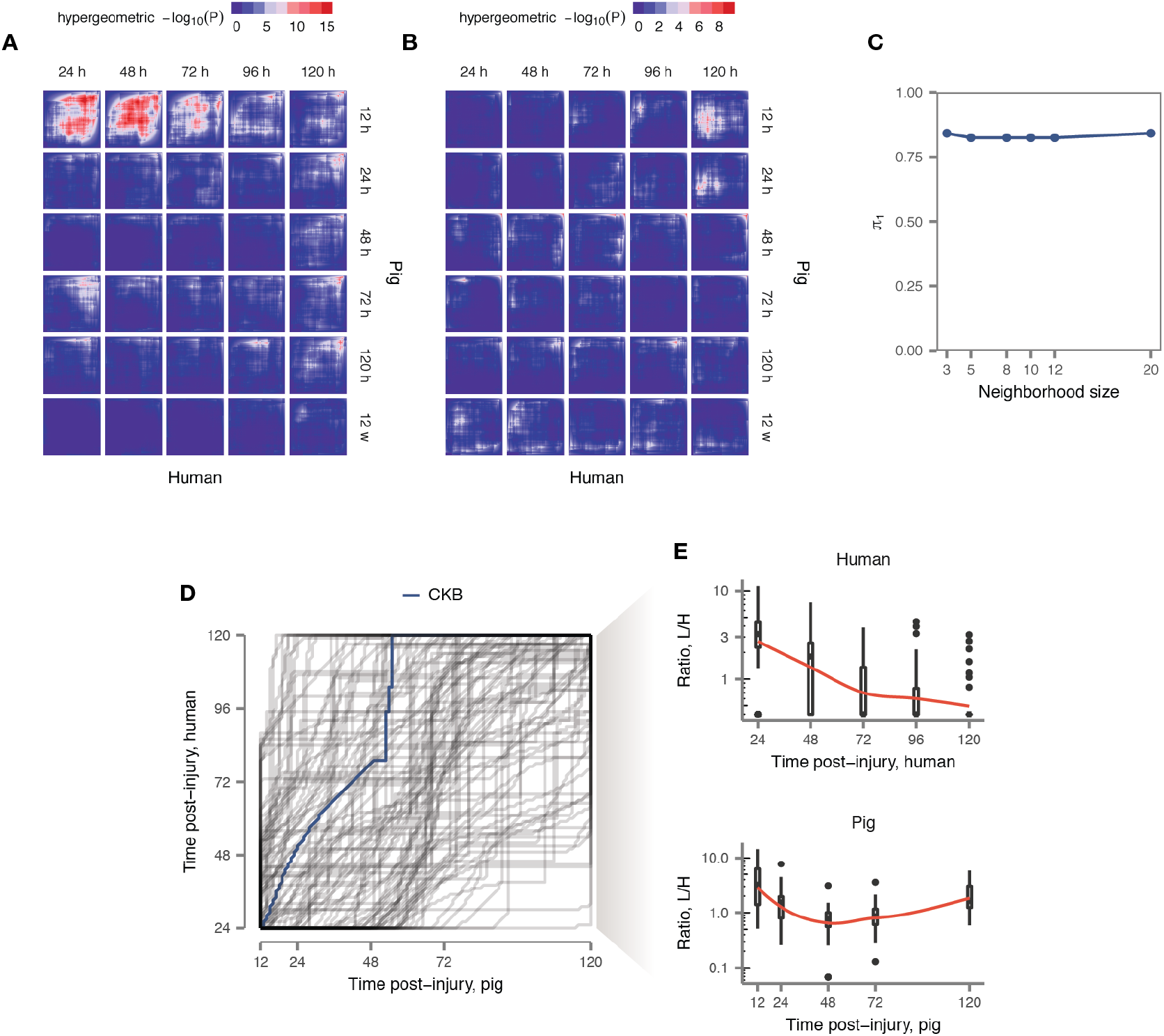
Cross-species conservation of the proteomic response to SCI. (A) Rank-rank hypergeometric overlap of proteins associated with injury severity at all pairs of timepoints post-injury. (B) Rank-rank hypergeometric overlap of proteins associated with neurological recovery at all pairs of timepoints post-injury. (C) Proportion of true associations, *π*_1_, estimated from neighborhood analysis of conserved coexpression p-value distributions with the neighborhood size varied between 3 and 20 neighbors. (D) Optimal alignments of individual proteins between the human and pig CSF proteomes over the first five days post-injury by dynamic time warping. CKB is shown in blue. (E) Abundance of CKB in human and pig CSF over the first five days post-injury. Red lines show local polynomial (loess) regression.

